# Associative memory of structured knowledge

**DOI:** 10.1101/2022.02.22.481380

**Authors:** Julia Steinberg, Haim Sompolinsky

## Abstract

A long standing challenge in biological and artificial intelligence is to understand how new knowledge can be constructed from known building blocks in a way that is amenable for computation by neuronal circuits. Here we focus on the task of storage and recall of structured knowledge in long-term memory. Specifically, we ask how recurrent neuronal networks can store and retrieve *multiple* knowledge structures. We model *each* structure as a set of binary relations between events and attributes (attributes may represent e.g., temporal order, spatial location, role in semantic structure), and map each structure to a distributed neuronal activity pattern using a vector symbolic architecture (VSA) scheme.

We then use associative memory plasticity rules to store the binarized patterns as fixed points in a recurrent network. By a combination of signal-to-noise analysis and numerical simulations, we demonstrate that our model allows for efficient storage of these knowledge structures, such that the memorized structures as well as their individual building blocks (e.g., events and attributes) can be subsequently retrieved from partial retrieving cues. We show that long-term memory of structured knowledge relies on a new principle of computation beyond the memory basins. Finally, we show that our model can be extended to store sequences of memories as single attractors.

---

Human memory is remarkable in its ability to robustly store and retrieve information with complex and hierarchical structure, guiding cognitive processes on many different timescales. In many instances, this “structured knowledge” can be described as sets of associations between discrete events with their contextual attributes. Some concrete examples are, temporal sequences representing events associated to particular times, episodic memories representing events associated with particular contexts^1,2^, cognitive maps representing spatial environments through landmarks associated with locations^3–5^, and semantic structures in language in which meaning is conveyed through sets of words associated with their respective roles within a sentence^6–8^.

To effectively use structured knowledge that has been stored in long-term memory, it must be represented in a way that allows for its retrieval through partial information, with tolerance for noisy and degraded cues. This is likely facilitated by the distributed nature of the underlying neural representations, which provide an inherent notion of similarity between representations and a mechanism for learning representations by the the tuning of synaptic weights in a neural network^7,9–11^. However, while the utility of distributed representations is clearly beneficial from this perspective, it is still not well understood how to represent associations and relations in neural networks in an efficient and flexible way that is amenable to the variety of computational demands involved in higher cognition^12–14^.

Several recent studies have addressed the contextual modulation of neuronal representations, e.g., by forming “mixed representations”^15^ or by gating parts of the network^16^. Other proposals have tried to implement more general relational structures in neural networks. An early attempt used the tensor product to create a distributed representation of pairwise relations between discrete items^6^. Subsequently, several Vector-Symbolic Architectures (VSA) were proposed as compressions of the tensor product to avoid the increase in dimensionality of the representation, allowing for the creation of hierarchies of relations in a compact way^17–22^. More recently, several architectures for deep or recurrent neural networks have been proposed to promote flexible relational reasoning^23–30^. However, these works have primarily focused on working memory, i.e., online tasks of processing incoming structured data. By contrast, the challenge of storing and retrieving relational structures in long-term memory has received little attention.

Storing knowledge structures in long-term memory poses several additional challenges. While working memory tasks typically process few structures at a time, long-term memory networks must cope with storing a very large number of structures, such as complex cognitive maps, multiple sequences, or stories, which may scale with the size of the memory network itself. While several works in the psychology literature^34^ Two generic measures of the efficiency of information storage in recurrent neural networks are their extensive capacity, i.e., the number of stored items scales with the number of neurons in the network^32^, and the ability to recall memories from partial cues which have small but significant overlap with the desired memory^33^. Both of these measures can be adversely affected by correlations across memorized patterns. For relational structures, additional correlations may occur due to the presence of objects or contextual attributes in multiple memories, putting additional constraints on the encoding of relational information. In addition to these considerations, models of distributed representation of knowledge structures typically compress the relational structure into a fixed-length distributed vector representation. To compensate for this loss of information, “clean-up” mechanisms are invoked. Thus, it is crucial that such mechanisms can be adapted for the task of retrieval of such structures from long-term memory to efficiently store large numbers of relational structures each containing multiple associations.

In this work, we propose a model for associative memory of *multiple* relational structures by using a quadratic binding scheme to form vector representations of memories consisting of *multiple* binary relations between items (which we will henceforth denote as pairs of objects and their attributes).

While our model is quite general, in most of our work we will use the holographic reduced representation (HRR)^7^ VSA scheme for convenience. We show that the binarized versions of these structures can be stored as *fixed point* attractors in a recurrent neural network and each structure can be retrieved from the memory network by using a cue which is a structure encoding a subset of the relations in the memorized structure. We highlight the holistic nature of this model by comparing the storage of temporal sequences in the present model, where the entire sequence is stored as a single fixed point, to previous models, where a sequence is stored as a sequence of transitions between multiple fixed points and cannot be fully recalled at once^35^. Our model posits that in addition to the network that stores the structures, a Dictionary network stores all individual items (e.g., individual words, familiar objects). We show that the identities of the objects contained in the structure can be decoded faithfully from the retrieved memory by querying the retrieved structure with the appropriate cue as long as a “clean-up” operation is performed to map the noisy estimate of the object to the correct item in the Dictionary. Furthermore, this decoding works well even when the retrieved structure is significantly degraded.

## Relational Structures

We begin by modeling a binary relational structure *S* as a set of *L* object/attributes pairs

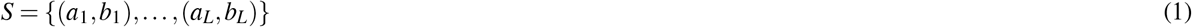

where both objects *a* and attributes *b* have embeddings as real vectors representing distributed patterns of activation in a neuronal population. For simplicity, both populations will be assumed to be of the same size *N*. We represent relations between items in a pair (*a, b*) by a transformation through a pairwise quadratic nonlinearity to a binding vector *g*(*a, b*) (in ℝ^*N*^) representing a distributed pattern of activity in a population of *N* “binding” neurons. Each component of the binding vector takes the form

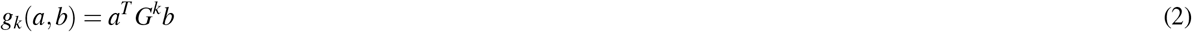

where each *G*^*k*^ is an *N* × *N* fixed binding matrix. The binding operation in Eqn. 2 is a generalized version of a VSA scheme^18^ and can be interpreted as a lossy compression of the tensor product binding operation first proposed in^6^.

We obtain the representation of the full relational structure *S* by the vector summation of the individual object/attribute pairs,

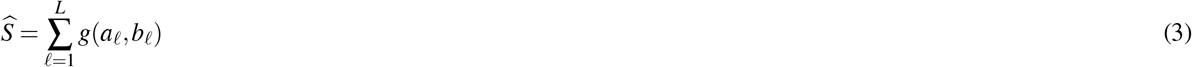

where the vector summation induces a second source of information loss. The representation 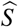 is permutation invariant with respect to the index ℓ so that the relations within the structure have no particular order.

The compressed representations of structures can be used for a variety of computations, such as structure classification. Here we focus on unbinding tasks, in which given 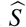 and one of its attributes *b*_ℓ_, we need to estimate its pair *a*_ℓ_. Similar to binding, we assume that the unbinding operation is performed through a quadratic transformation involving the pair 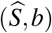, so that the *k*-th component of the estimator 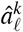 of *a*_ℓ_ is given by

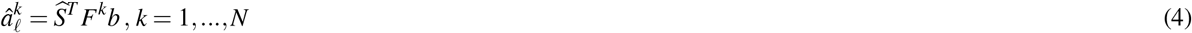

where each *F*^*k*^ is an *N* × *N* matrix chosen so that the decoding operation is the approximate inverse of the binding operation.

In general, the binding and unbinding matrices can be learned and the optimal choice should depend on the nature of the items contained in the dictionary. Here we use a generic set of matrices, a popular choice known as Holographic Reduced Representations (HRR) described in Methods.

The final estimate of *a, ã*, is computed by comparing the noisy estimate against a Dictionary, i.e., a neural long-term memory system that stores all familiar objects *a*_*d*_, using,

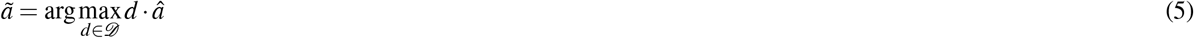

A schematic of the encoding and decoding networks is shown in Fig. 1a.

**Figure 1.**
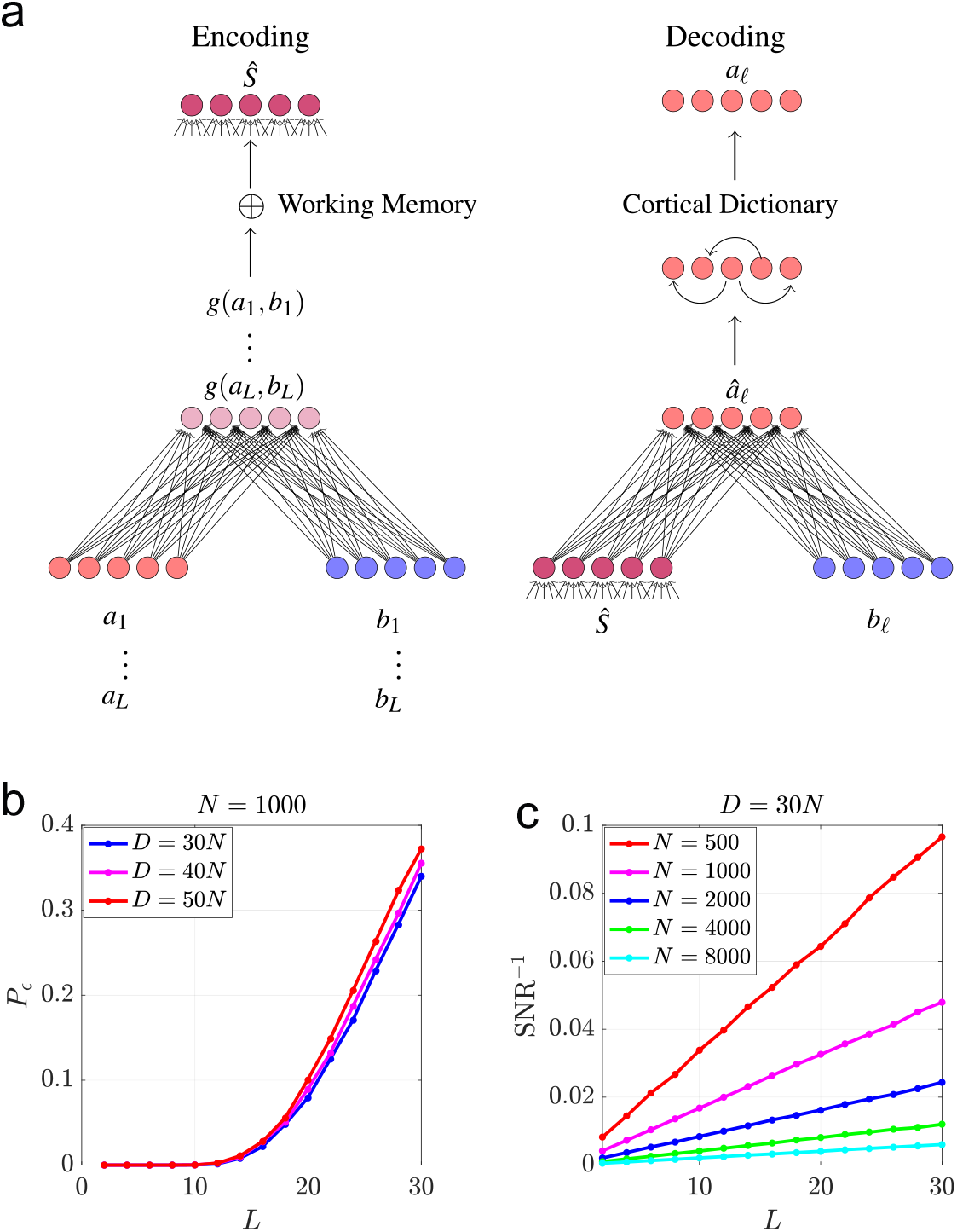
(a) A schematic of the network used to bind objects/attribute pairs (*a, b*) to form the knowledge structure 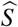 alongside the network used to decode an object *a*_ℓ_ from 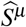 by presenting the attribute *b*_ℓ_. (b) The unbinding error *P*_*ε*_ shown as a function of structure length *L* for *N* = 1000 for various values of *D* averaged 10, 000 structures (c) SNR^−1^ shown as a function of *L* for several values of *N* averaged over 10, 000 structures.

The maximum likelihood (ML) decoding error is given by the probability *P*_*ε*_ that the estimator *â* has the largest overlap with an incorrect item in the dictionary. It depends on the size of the dictionary *D* and the signal-to-noise ratio (SNR) overlaps of the estimator *â* with items dictionary defined in Eqn. 20 in Methods. ML decoding is an idealization of a more biologically realistic retrieval of the stored pattern *a* from a retrieval cue *â* in a long-term associative memory network storing all individual Dictionary items. Two possible implementations of this memory system are a winner take all network with lateral inhibition^36^ or a sparse Hopfield network^37^.

### Storing Structures in Long-Term Associative Memory

We now consider the long-term memorization of multiple knowledge structures by storing their vector representations in a neural network for long-term Structured Knowledge Associative Memory (SKAM), so that they can be retrieved at a later time from partial cues and subsequently queried to reconstruct individual events. We consider a set of *P* structures {*S*^1^, …, *S*^*P*^} which for simplicity, all consist of *L* items. We label the set of L objects and attributes comprising the *μ*-th structure as 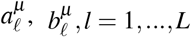.

For each structure, the HRR encoding scheme is used to create a vector representation 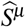 from *S*^*μ*^.

To store multiple structures as fixed points in a neural network, the neuronal input-output transformation must be highly non-linear, implying that the stored patterns themselves are limited to the dynamic range of the neurons. As in a standard Hopfield network^32,33,38^, we assume neurons are binary ±1 variables and the memory patterns, candidates of fixed points of the attractor dynamics are 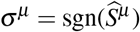. For a network with *N* binary neurons, the memory load is defined as 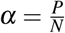, where *P* is the number of stored structures.

In general, the associative nature of Hopfield memory networks is expressed in the ability to recall a memorized pattern starting in any initial state which has a sufficiently large overlap with the memorized pattern. If the initial state is within the basin of attraction of the pattern, it will converge to the pattern without errors. In our case, we consider partial cues of a structure, 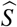, in the form of a recalling structure 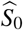 obtained by the binding and subsequent summation of any subset *S*_0_ of the *L* pairs of binary relations contained in 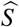, i.e.,

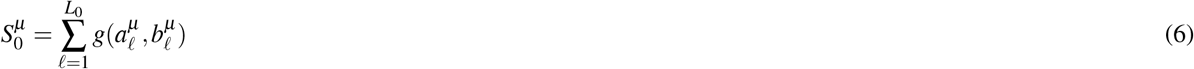

where *L*_0_ is the number of recalling elements in 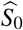 and is assumed to be much less than *L*. The network is then initialized in the state 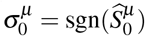 and evolved to a fixed point which, if successful, corresponds to the stored binarized structure *σ*^*μ*^. A schematic of our model is shown in Fig. 2 and more details on the memory network are given in Methods.

**Figure 2.**
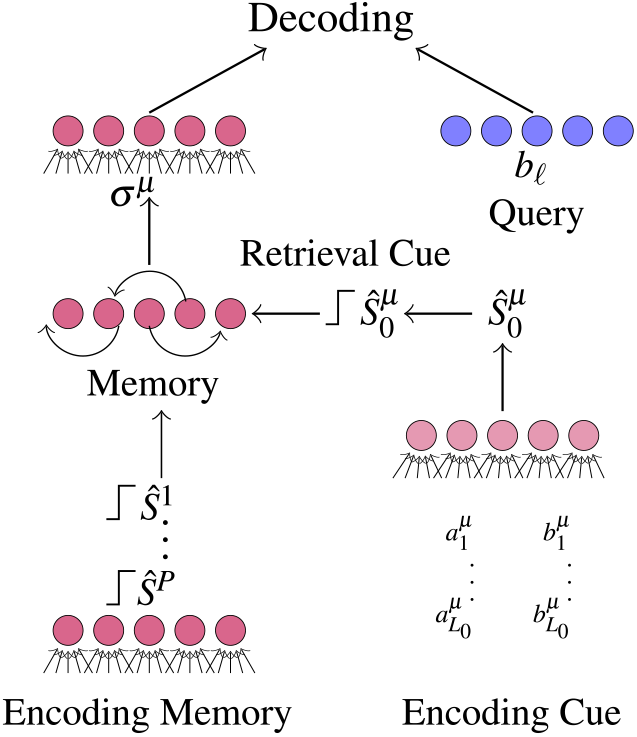
A schematic of the process of storing multiple binarized structures in a memory network. These structures can be retrieved from memory by encoding a retrieval cue from a subset of the relations in the desired structure as in Eqn. 6 and initializing the network in the state of the binarized cue.

There are several learning rules which can be used to store patterns as discrete fixed points in recurrent neural network models of associative memory. Here, for simplicity, we use the Pseudo-inverse learning rule proposed in^39^ to train the network. In a Pseudo-inverse network, all structures are perfect fixed points for *α* < 1, which is assumed throughout. This allows us to focus on the retrieval cues, since failure to perfectly recall a structure occurs only when the retrieval cue is outside of the basin of attraction of the memorized structure. We observe qualitatively similar behavior for the Hebb learning rule and the Storkey rule introduced in^**?**^ for *α* well below the memory capacity described in Section 3.4 of the Supplementary Material.

## Results

We evaluate the performance of the scheme introduced in the previous section by the ability to accurately perform the unbinding operation after retrieval of a structure from the SKAM. For example, after retrieving the structure *μ* = 1, we should be able to extract the item 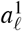 with a query in the form of its pair, i.e., 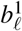 with low error. We quantify performance by the average unbinding error *P*_*ε*_ obtained in simulations where structured memories are created from random patterns, stored in memory, retrieved with partial cues 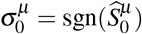, and subsequently decoded using the ML “clean-up” operation. We assume all items appearing in memorized structures are stored in dictionaries for objects and for attributes which are then used to decode from the retrieved memory. A schematic of this process is shown in Fig. 2 and full details of the simulations are provided in Methods.

The parameters involved in the performance measure are: network size *N*, memory load *α* = *P/N*, size of the relational structures and the retrieval cue, denoted as *L* and *L*_0_, respectively. We consider the regime where both *N* and *P* are very large and the memory load *α* ∼ *O*(1)^32,40^, mainly considering values *α* ∼ 0.1 − 0.2, where the network acts as a good associative memory.

### Retrieval of Structured Memories

We begin by showing numerical results which measure the quality of the retrieved structures in terms of the unbinding error *P*_*ε*_ and the SNR of overlaps defined in Eqn. 20. In all reported results, the extracted item (and the associated query) comes from pairs that are not part of the cueing structure *S*_0_. Thus any performance better than chance necessarily involves information extracted by the retrieval from long-term memory. In Fig. 3a we show the dependence of the unbinding error *P*_*ε*_ on *L, L*_0_, and *α*. For comparison, we show *P*_*ε*_ for the original structure prior to storage in the memory network, demonstrating that except for small *L*, the dominant contribution to the error comes from retrieving the structure from long-term memory. We also observe that for a fixed *L, L*_0_, and *α*, the error is suppressed as *N* increases, in contrast to standard large attractor memory networks where performance depends only on *P/N*. To elucidate this behavior, we replot the results in terms of the SNR^−1^, i.e., the inverse of the SNR as defined above in Fig. 3b, showing that for each *L*_0_, there is a critical *L* above which the SNR of the memorized structures decreases relative to the original SNR, SNR_0_ before storage in long-term memory. Note that due to binarization, SNR_0_ is smaller by a factor of 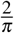 relative to the value given in Eqn. 22. we replot the same results in terms of the inverse of the normalized SNR, SNR*/*SNR_0_ vs., *L/L*_0_. Since SNR_0_ is proportional to *N/L*, this normalization factors out the “trivial” dependence on *L/N* from the post retrieval SNR. Figure 3c shows that for a fixed *α* the normalized inverse SNR depends only on *L/L*_0_. and only weakly on *N*, suggesting that the main *N* dependence comes from the linearity of SNR_0_ in *N*.

**Figure 3.**
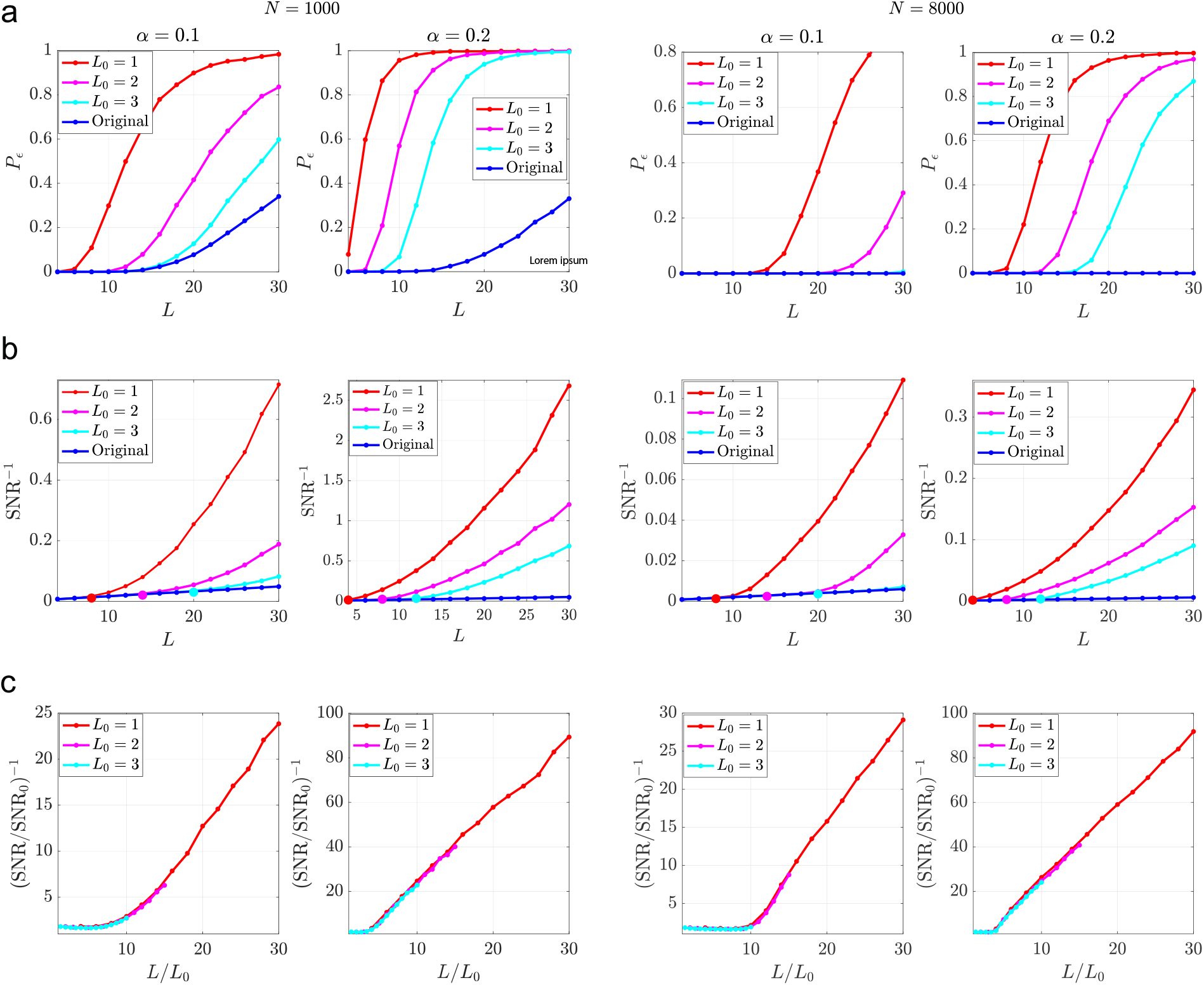
(a) The decoding error *P*_*ε*_ is compared for several values of *L*_0_ from structures containing *L* pairs of items for two values of *N* and two values of *α*. (b) SNR^−1^ v. *L*. For each value of *L*_0_, *l*_*c*_ is given by the value of *L* where SNR^−1^ diverges from the line corresponding to the original structure, which is marked for each value of *L*_0_. (c) (SNR*/*SNR_0_)^−1^ v. 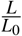 where 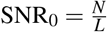. For all figures, *T* = 20 parallel updates are used in memory retrieval and the average is performed over 10, 000 memories. The dictionary is fixed to *D* = 30*N*.

### Length of Cueing Structure and Memory Basins

As seen in Fig. 3, the performance worsens (and SNR decreases) as *L* increases, while the converse holds true for *L*_0_. We find that there is a critical ratio, 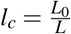, defined as the minimum initial cue (relative to the total length), that leads to very small error which is essentially equivalent to the error for the original structure.

To understand the origin of *l*_*c*_, we note that 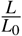 specifies the average initial overlap of the retrieval cue with the corresponding memorized structure, which we denote *m*_0_. For small values of 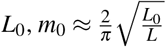. The size of *m*_0_ determines whether on average the initial state is within the basin of the desired memory, so that the recurrent dynamics will succeed (or fail) in converging to the desired attractor. As the cue length *L*_0_ grows, the initial state becomes increasingly likely to be within the basin of attraction of the desired structure, retrieving it with essentially no error. In these conditions, the unbinding operation has the same probability of success as for the original structure. Conversely, for small enough *L*_0_ the initial state is likely outside the attraction basin of the memory, leading to errors in the retrieved structure.

To determine the minimum value of *L*_0_ required for perfect retrieval, we use known estimates^39^ of the radius of attraction in attractor memory networks, *R*(*α*) = 1 − *m*_*min*_(*α*), where *m*_*min*_(*α*) is the minimal overlap between the initial state and the desired memory required for convergence to the correct fixed point on average. *m*_*min*_(*α*) determines the minimal length of the cueing structure, i.e., 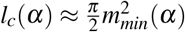.

We conclude that when *L*_0_*/L* < *l*_*c*_, the main source of the decoding error in our model comes from the limitation on good retrieval of the structure from memory, due to small values of *L*_0_*/L*, and not from noise in the original encoding (corresponding to *L*_0_*/L* > *l*_*c*_).

### Retrieval Outside Memory Basins

Naively, one would expect that for *L*_0_ < *l*_*c*_*L, P*_*ε*_ will be very large due to the accumulation of errors in the retrieved structure, which is outside the memory basin. However, as shown in Fig. 3, this is not case. Surprisingly, the decoding performance is well below chance level for substantial range of values of *L*, even when *L* ≫ *L*_0_*/l*_*c*_. This observation can be explained by two scenarios: (1) the actual basins fluctuate in their shape so that for some structured memories, initial states may converge to the memory fixed point even if they are outside the *mean* basin radius; (2) initial states outside the true memory basin converge to fixed points with significant overlap with the structured memory.

To test these scenarios, we measured the empirical distributions *p*(*m*) where *m* is the overlap between the fixed point and the desired structure, obtained from histograms of overlaps for several values of 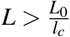, shown in Supplementary Fig. 3.

We find that as *N* is increased, *p*(*m*) becomes sharply peaked around a single value *m*\*. Inside the basin of attraction, i.e., *m*_0_ < *m*_*min*_(*α*), *m*\* = 1. However, outside of the basin when *m*_0_ < *m*_*min*_(*α*), *m*\* < 1; nevertheless it is substantially larger than 0. The value of *m*\* depends on both *m*_0_ and the load *α* roughly as

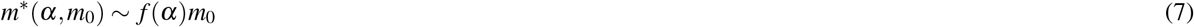

described in further detail in Methods and the Supplementary Material. 3. A schematic of the energy landscape is shown in Fig. 4a. Furthermore, for *α* ≲ 0.3, *f* (*α*) > 1 (Fig. 4b), implying that the final overlap with the retrieved structure is significantly larger than the initial overlap *m*_0_ even far outside the basin of the structure.

**Figure 4.**
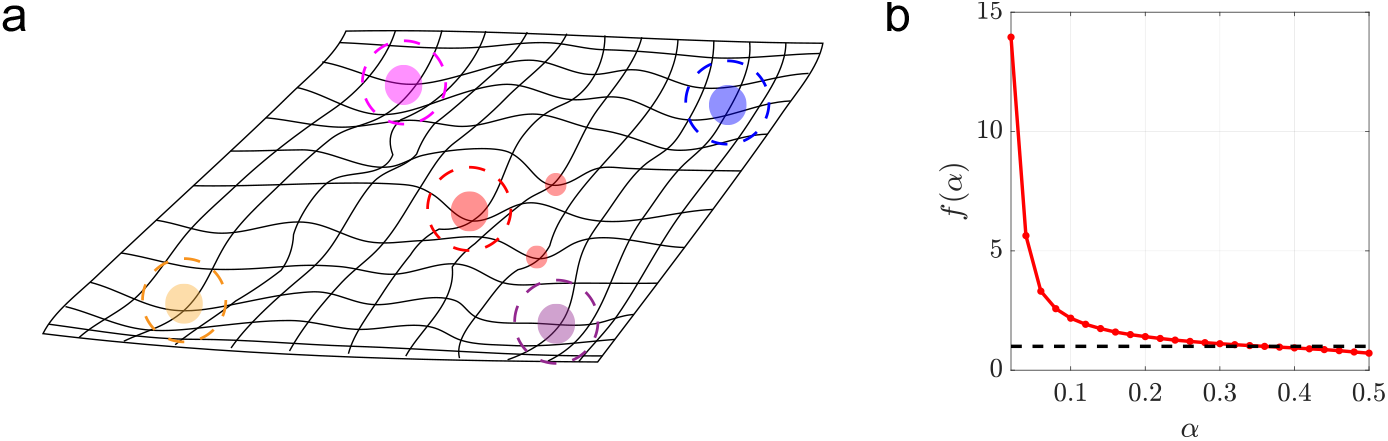
(a) schematic of the energy landscape of the memory network. The large filled circles are stored memories and the dashed circles denote their basins of attraction. The two small red circles are fixed points outside of the basin of attraction of the red memory which still lead to a large enough overlap for accurate decoding when 1 ≫ *N/L*. (b) *f* (*α*) defined in Eqn. 7 is obtained from simulations of a Pseudo-inverse network storing random memories of size *N* = 8000. The black dotted line at 1 shows that *m*\* > *m*_0_ for *α* ≲ 0.3. *T* = 20 parallel updates are used for memory retrieval and averages are performed over 50 trials.

### SNR of Retrieved Structures Outside the Basin

We use the preceding results to estimate the SNR for *L*_0_*/L* ≪ *l*_*c*_, i.e., when the initial state is well outside the memory basin. First, we argue that the SNR of unbinding from a noisy state with overlap *m* < 1 with the true structure, should be roughly,

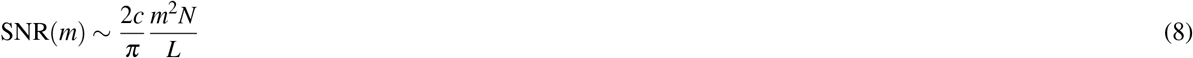

where, as before, the factor of 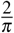 comes from binarization and *c* ≈ 0.65 accounts for the fact that part of the overlap *m* is contributed by the initial cueing structure *S*_0_ and is more concentrated around the relations contained in the retrieval cue. For very large networks, we can replace *m* in Eqn. 8 with *m*\* from Eqn. 7. Using Eqn. 30 from Methods, we express *m*_0_ in terms of *L*_0_*/L* and arrive at

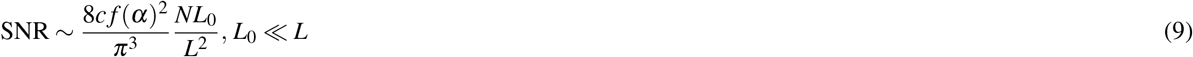

which is verified in Fig. 5 for two values of *α*. These results summarize the rich behavior of associative memory of structured knowledge. In contrast to standard memory functions, here the performance depends not only on the memory *α* but also on the network size *N*, structure length *L*, and cueing length *L*_0_ through the SNR. The key difference is that in structured memories, the criterion for success is not limited to convergence to the target memory; even if the target memory is only partially retrieved, the underlying memorized relations can be still be retrieved faithfully using the semantic memory. The well-known property of pattern completion is realized here by a sub-structure of length *L* < *L*_0_, in addition to the standard random initial condition.

**Figure 5.**
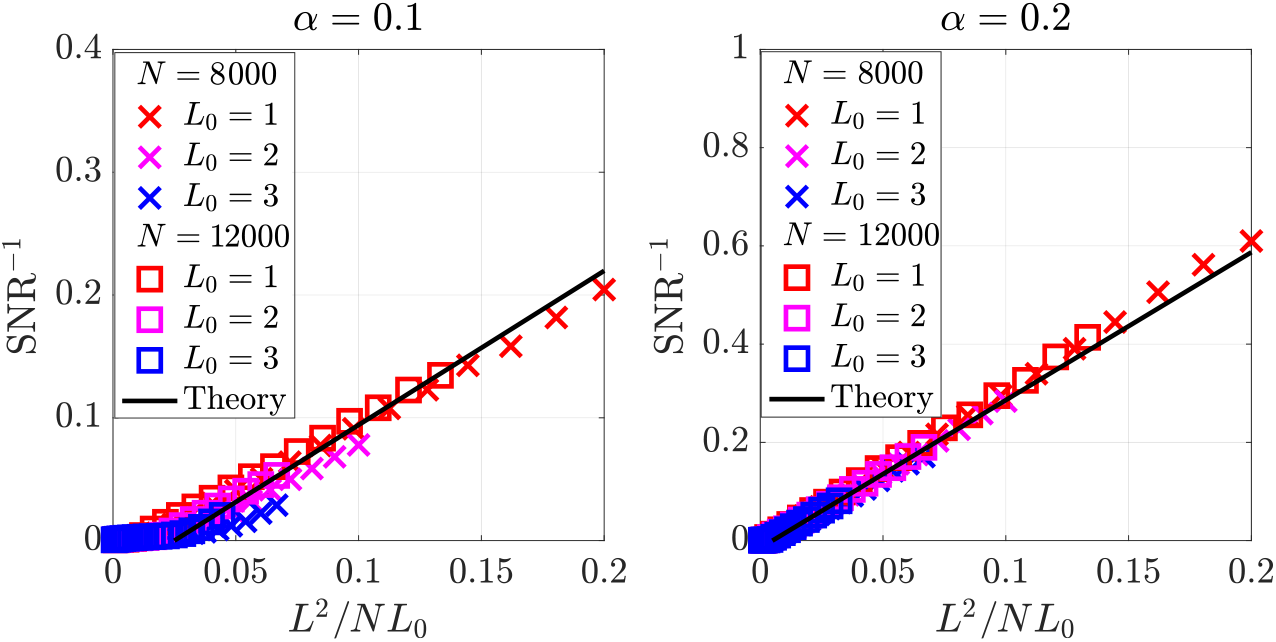
SNR^−1^v. *L*^2^*/NL*_0_ shown for *N* = 8000 and *N* = 12000 and for several values *L*_0_ as an average over items contained in 10, 000 memories shown for *α* = 0.1 and *α* = 0.2. The black line is obtained from Eqn. 9 with *c* = 0.65. *T* = 20 parallel updates are used for memory.

### Storage and Retrieval of Sequences

#### Storing Sequences as Binary Structures

We now extend the results of the previous sections to representations of temporal sequences. Temporal sequences can be modeled as structures in several ways. One possibility is to bind each event in the sequence with its temporal order in the sequence. This can be implemented via a contextual drift process with a context representation that evolves as items in the sequence are retrieved as in the Temporal Context Model of free recall of lists^2^ and similarly in the Context Retrieval and Updating model^41,42^. Here we use temporal proximity as the contextual cue by interpreting a sequence as a set of binary associations between temporally proximal events. Thus, a temporal sequence of length *L*, (*a*_1_, *a*_2_, …, *a*_*L*_) can be represented as a structure of the form

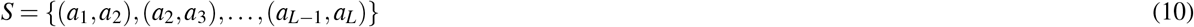

and the entire sequence *S* is represented by a vector 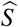 of size *N* given by

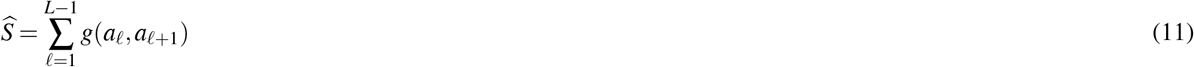

Decoding an episode at a particular time, i.e., *a*_ℓ_, is performed through an unbinding operation with a query by the preceding event, *a*_ℓ−1_. Starting from a query by *a*_1_, the entire sequence can be unfolded through a sequence of queries. Because each event appears in two binary relations, we need to use an asymmetric binding operation so that *g*(*a, b*) ≠ *g*(*b, a*). Within HRR, this can be accomplished by switching the binding and unbinding operations^43^.

As before, we consider the case in which all items being decoded are contained in a Dictionary 𝒟 = {*a*_1_, *a*_2_, …, *a*_*D*_}, so each decoding step involves a clean-up of the decoded item before preceding to decode the next item from the sequence. A schematic of this process is shown in Fig. 6a.

**Figure 6.**
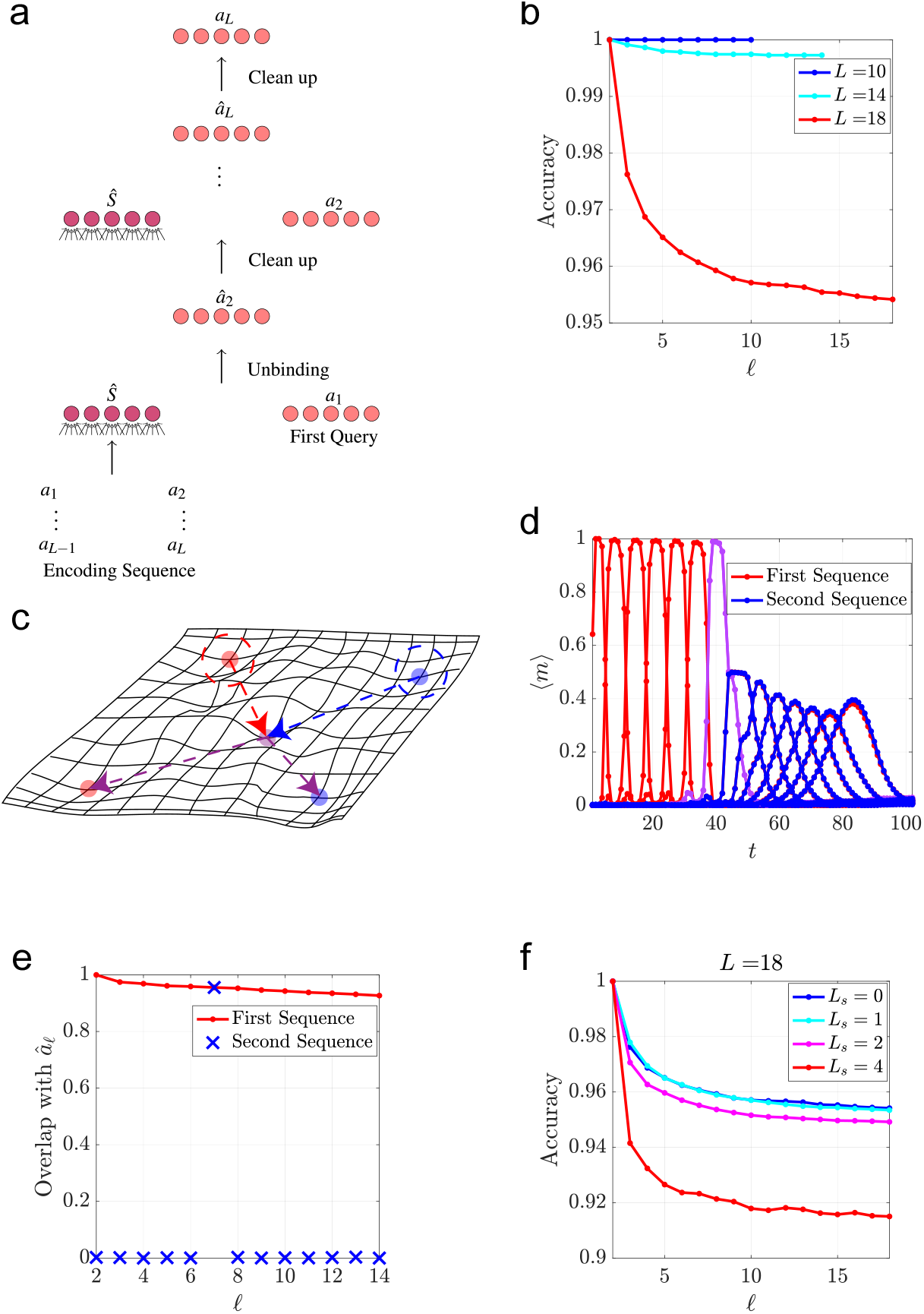
(a) A schematic of the decoding scheme for sequences. (b) Decoding accuracy as a function of sequence position ℓ along sequences of length *L* retrieved from memory shown for several values of *L. N* = 2500, *α* = 0.05, and the average is computed over 100 trials. (c) A schematic of the energy landscape for two temporal sequences containing a single common element which are encoded as sequences of attractors using the scheme in^35^ (discussed in Section 4 of the Supplementary Material). (e) The average overlap of the network state ⟨*m*⟩with each attractor in the sequence is shown as a function of retrieval time for a network of size *N* = 1000 storing two sequences containing a single element in common. The parameters *τ* = 8 and *λ* = 2.5 are used and the average is computed over 1000 trials. (e) The average overlap of the estimator *â*_ℓ_ (normalized by ⟨ *â*_2_ · *a*_2_ ⟩) with the correct Dictionary item at each position in the sequence for a network of size *N* = 1000 storing two sequences containing a single element in common. The average is computed over 10, 000 trials. (f) Decoding accuracy as a function of sequence position ℓ for *P* = *αN* sequences for which each neighboring sequence contain *L*_*s*_ common elements with the previous one. *N* = 2500, *α* = 0.05 and the average is computed over 100 trials.

The binarized versions of the structures representing each sequence are stored for long-term memory in a recurrent neural network with synaptic weight matrix determined via the Pseudo-inverse rule. The cueing structure 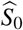 consists of the first relation (*a*_1_, *a*_2_), so the overlap of 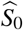 with the stored sequence 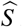 is closely approximated by Eqn. 30 with *L*_0_ = 2. Alternatively, the first item *a*_1_ can be used as a retrieval cue if it is added to the representation 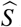 in Eqn. 11. We are primarily interested in the ability to reconstruct the entire sequence after it is retrieved from memory.

#### Retrieval of Sequences from Long-term Memory

Due to the sequential nature of decoding sequences in our model, the decoding error accumulates as each subsequent element is retrieved. Thus, the unbinding error for an event depends on its position in the sequence (relative to the cued events). In Fig. 6b, we show the decoding error *P*_*ε*_ at each position along sequences encoded by Eqn. 11 for sequences of different length *L*. Since the SNR of the overlap with the correct item along each position in the sequence depends on *L*, the length of the sequence limits the accuracy of decoding at all positions along the sequence. Nevertheless, for low memory loads and moderately long sequences, the accumulated error is small.

#### Decoding Error

An interesting outcome of this mode of recall is the accumulation of errors as the recall sequence advances. This would give rise to correlations between probabilities of recall that decay as a function of the temporal lag between the events, consistent with observations^2^. This behavior was previously explained by positing that proximal temporal context vectors are correlated. In our model, these correlations are a natural consequence of the fact that the proximal events serve as temporal context cues.

The present scheme of long-term storage of sequences as single fixed points overcomes a key disadvantage of previous attempts at storing multiple temporal associations in attractor neural networks. In previous models of sequential memory^35^, all of the patterns contained in the sequences are stored as separate attractors in one network and the sequences themselves are encoded in a time-delayed synaptic plasticity rule that associates each pattern with its next pattern in the sequence, illustrated in Fig. 6c and reviewed in Section 4 of the Supplementary Material. Because of the Markovian nature of the synaptic plasticity, retrieval will fail if multiple sequences share the same event, as shown in Fig. 6d. By contrast, in the present model, the entire sequences are stored and retrieved as separate “holistic” attractors. Thus, as long as the retrieval cue is unique to a single sequence, it will retrieve it unambiguously and the subsequent unfolding of the sequence by the unbinding network will be immune from interference with other sequences. To demonstrate this, we consider *P* sequences *S*^*μ*^ with *μ* = 1, …, *P* of length *L* where neighboring sequences *S*^*μ*^ and *S*^*μ*+1^ share *L*_*s*_ events in common.

In Fig. 6e, we show the decoding error for a sequence stored in memory with another sequence containing an overlapping event, demonstrating the successful retrieval of the entire sequence despite the presence of an overlapping state (compare with Fig. 6d).

Figure 6f shows that sequences can be faithfully retrieved even with multiple common states, up to the point where the basins of attraction of individual sequences shrink due to the large overlap between them.

Finally, it is interesting to compare the the memory capacity of the two models. In the sequence attractor model, the maximal number of stored sequences of length *L* is *P* < *α*_*c*_*N/L* since the network stores *PL* states. In contrast, in the present model, since only *P* attractors are stored, the capacity of storage is *P* < *α*_*c*_*N*. Nevertheless, for a successful unfolding of the sequence we need *PL/N* to be bounded (for a fixed *L*_0_*/L*) due to the noise in the unbinding operation. A potential disadvantage of the current model is the need to devote additional memory resources to store the individual events in a Dictionary. On the other hand, the Dictionary can be used for multiple other cognitive tasks aside from the retrieval of these sequences. Another flexibility in the separation of the retrieval of the neural representation of the sequence from the subsequent reconstruction of individual events, is the fact that, for some tasks, the agent may not need to access the full detail of the sequence, for instance in tasks that requires distinguishing between one episode and another one. Such tasks may not need to rely on the full unbinding of the sequences.

### Neural Implementation of Multiplicative Binding

We now briefly consider possible implementations of the binding computation in Eqn. 2 through multiplicative interactions in biological neurons^44^. Previously, several mechanisms have been proposed to facilitate multiplicative interactions among neurons including dendritic gating^45^, quadratic firing rate response^46^, and short-term synaptic plasticity. Short-term plasticity comprises a variety of synaptic processes that modulate synaptic efficacy in a transient, activity-dependent manner^47^. These processes occur on timescales ranging from milliseconds to minutes^48^ and are thought to mediate higher cognitive functions like attention and working memory. More recently it has been suggested that “fast weights” in artificial neural networks may serve as an analogy to short-term plasticity in the brain^49^ with connections to linear transformers^50^.

We start by noting that *g*_*k*_(*a, b*) = *a*^*T*^ *b*’ where *b*’ = *G*^*k*^*b*. The last term is a representation of the activity pattern *b* by propagating it through a synaptic matrix *G*^*k*^. Finally the dot product between *a* and *b*’ can be implemented can be decomposed into an outer product of two fixed synaptic weight vectors, i.e. 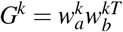 so that components of the binding vector take the form

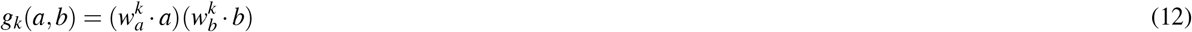

We now use the above form to consider how firing rate nonlinearity and short-term synaptic plasticity can serve as mechanism for generating quadratic binding.

#### Nonlinearity of the Firing Rate

Biological neurons can potentially implement the computation of the binding vector *g*(*a, b*) via the nonlinearity of the firing rate response to a synaptic current *r* = *f* (*I*) where the synaptic current is given by the sum 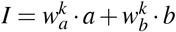. While for many neurons rectificed nonlinearity *f* (*I*) = [*I* − *I*_0_]_+_ is a good approximations, other neurons are found to be approximated by quadratic nonlinearity 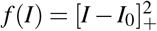 where the firing in response to the sum of the separate responses to each input *a* and *b* is subtracted from the response to the combined input from *a* and *b*. Building on a quadratic *f* (*I*) curve, we can write

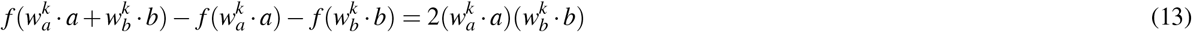

The subtraction can be implemented by inhibitory neurons or by temporal derivative in a working memory system.

#### Short-term Synaptic Plasticity

Another potential mechanism to generate quadratic binding is short-term synaptic plasticity. This can be accomplished by a short-term increase in residual presynaptic calcium levels in working memory enabling *b* to modulate the synapses *G*^*k*^ so that subsequent input *a*^*T*^ will generate the postsynaptic potential

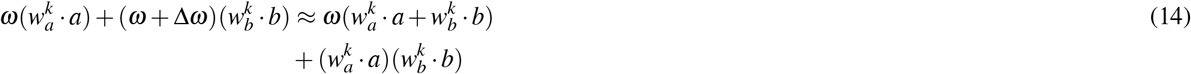

which contains a multiplicative component of the form in Eqn. 12. Note that Eqn. 14 contains a linear term weighted by ***ω***. This term may not need to completely cancel as it provides the trace with some similarity to both *a* and *b*, potentially allowing objects or context to be independently used as a retrieval cue. However, if *ω* relatively small, the trace will remain most similar to the bound conjunction *g*(*a, b*).

## Discussion

In summary, we have proposed and analyzed a model demonstrating how multiple knowledge structures, i.e., sets of relations between pairs of items, can be represented, stored, and retrieved in Hopfield type recurrent neural network models of long-term memory. Our model hypothesizes that the entire set of relations is encoded through binding operations, summation and binarization, in a single pattern of activity in a neuronal population, which is then stored as fixed point in the recurrent network. Retrieval of relational information from long-term memory, consists in our model of two stages: first, retrieval of the desired fixed point, and subsequent unbinding to uncover individual relations with the aid of a separate memory system, the Dictionary. Our analysis of this model clearly shows that the decoding SNR exhibits the appropriate scaling of parameters required for accurate decoding of objects coexisting among many other relations within a structure, and also among the extensive number of other structures stored in memory.

We also show that this scheme can be used to model long-term memory of temporal sequences by creating structure vectors for sequences of temporally associated items and store in a recurrent network compressed versions of the sequences as fixed points. Sequence recall consists of retrieval of the “sequence” fixed point, and subsequent unfolding of the stored events through a sequence of unbinding operations. In this application we have also demonstrated that our model for storing structure vectors in long-term memory is not very sensitive to the presence of a partial overlap between different structures.

Our analysis suggests that the success of this long-term memory system depends not only on the memory capacity of the attractor network but also very crucially on the properties of the memory basins of attraction and the landscape in the surrounding “terrain”, such as the degree of overlap between “spurious” states outside the basins with the target fixed point (inside it). For this reason, a learning rule that decorrelates memories and yields smoother basins is clearly superior, as shown in Supplementary Fig. 8. Due to the dense distributed nature of the binding scheme employed here (HRR), we have not studied the effect of pattern sparsity on the long-term memory system^37^. It would be interesting to explore the sparsity effect in sparse binding schemes^51–54^ and generally how the binding matrices can be learned in a biologically plausible way.

We close by briefly discussing two important aspects of this work which have the most immediate phenomenological implications. A key aspect of our model is the existence of neuronal populations representing entire relational structures in long-term memory as persistent patterns of activity displaying the “holistic context” of each structure. This system interacts with a working memory system which executes the dynamics of retrieving details of the stored relations. We have not addressed the interesting question of the mechanism by which a stream of experiences is segmented into a sequence of discrete events^55^, or more generally, the mechanism that segments complex environments into a discrete sets of bound items and how these representations may evolve over time^56,57^. In particular, our model of long-term memory of sequences predicts that the retrieval of a temporal sequence is associated with a persistent pattern of activity (representing the context of the entire sequence) in addition to sequential dynamics involving the dynamic interaction between working and long-term memory. This can be tested in recordings of neuronal activity during recall of sequences in the hippocampus and in songbirds. It would also be interesting to see how this fits in with studies of the dynamics of recognition memory in the psychology literature^56,58^.

Finally, as mentioned above, our framework of storage and retrieval of relational knowledge structures in long-term memory relies on the existence of a complementary long-term memory system, the “Dictionary”, which stores the individual building blocks comprising the relational knowledge. It is tempting to identify these two complementary memory systems as representing episodic memory (the relational system)^1^ and semantic memory (the Dictionary)^59^, although we emphasize that in the present context, semantic memory does not necessarily require language and presumably exists in other species as well. The synergy of these two “complementary” memory systems results in an associative memory system with both the capacity and flexibility to store and faithfully represent complex knowledge structures in long-term memory in analogy with the “complementary learning systems” framework proposed in^60^ and revisited in^**?**,61^. Adapting this framework to further explain empirically observed phenomena in memory will require adherence to known biological properties of hippocampal representations as well more explicit models of both the Dictionary and the working memory system in which binding and unbinding occurs.

## Methods

### Holographic Reduced Representation

HRR^8,62^ is a commonly used VSA scheme with fixed forms for the binding and unbinding matrices in Eqns. 2 and 4. The binding operation *g* is given by the circular convolution operation of the vectors *a* and *b* where

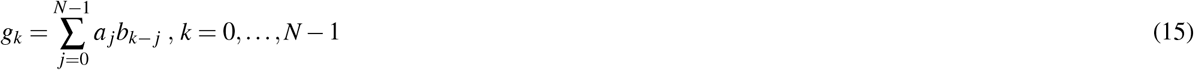

and all subscripts are defined modulo *N*. The circular convolution operation is both associative and commutative. The corresponding decoding operation *ϕ* is realized through circular correlation of the two vectors 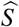 and *b* where

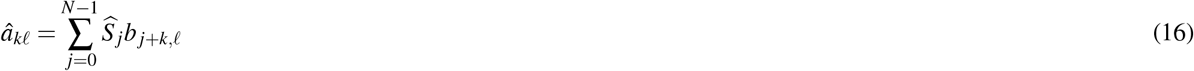

We see from comparing Eqns. 2 and 4 with Eqns. 15 and 16 that HRR corresponds to the following choice for the encoding and decoding matrices

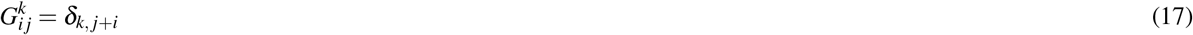

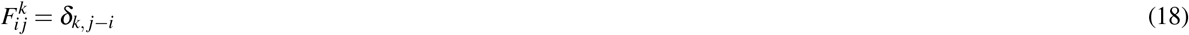

The commutativity of the encoding operation implies that HRR encoding is commutative. To represent non-commutative asymmetric relations, we can simply exchange the binding and unbinding operations i.e., binding with circular correlation and unbinding with circular convolution^43^. The full details of the statistics of decoding for HRR is given in Section 1.2 of the Supplementary Material.

#### Unbinding Accuracy

We access the typical decoding performance by considering the case in which *a*’s and *b*’s are random vectors with components drawn iid from 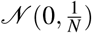 and the dictionaries for *a*’s contains *D* elements. Then the ML decoding error is well approximated by

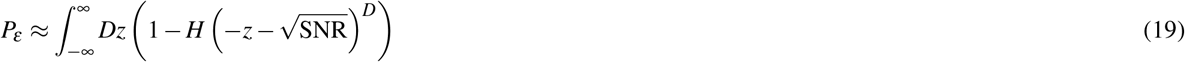

where 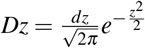 and 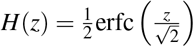. Here SNR is a signal-to-noise-ratio defined in terms of the mean overlap of the estimator *â*_*d*_ with the correct Dictionary item *a*_*d*_ and the variance of the overlap with incorrect Dictionary item *a*_*d*_’, i.e.,

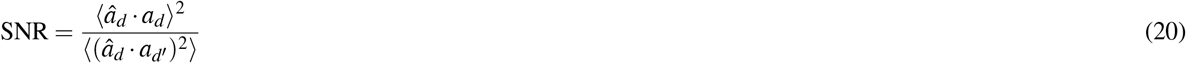

where *d*’ ≠ *d*, and the average is over the Gaussian distributions of the components of *a*_*d*_ and *a*_*d*_’. For full details see Section 2 of the Supplementary Material. For SNR ≫ 1, the decoding error can be approximated as

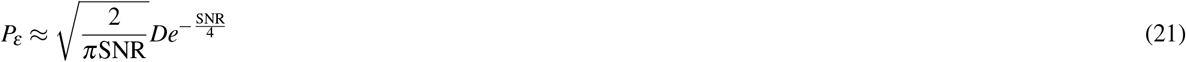

To leading order, the SNR for many VSA binding schemes (including HRR) is

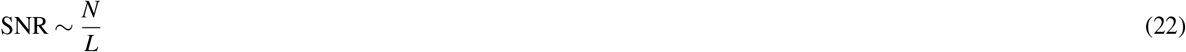

Eqn. 21 implies that *P*_*ε*_ ≪ 1 as long as *N* ≳ *O*(*L* log *D*). Hence, for *L* ≪ *N* accurate decoding requires the size of the Dictionary *D* be at most polynomial in *N*. In this regime, assumed throughout, the size of the Dictionary has little effect on performance, which is dominated by the SNR. *P*_*ε*_ and the inverse SNR are shown as functions of *L* in Fig. 1b and c, respectively.

#### Memory Network

Throughout this work, we consider Hopfield type recurrent neural networks with binary neurons. The state of the network at time *t, σ* (*t*) is given by the update rule

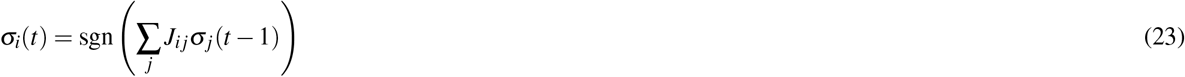

where updates are done either in series or in parallel. For simplicity, parallel updates are used for the figures in the main text, but we show in Section 3.3 of the Supplementary Material that the results are qualitatively similar for serial updates.

Given a set of memories *σ*^*μ*^, the synaptic weight matrix *J*_*i j*_ must be chosen so that each of the memories is a fixed point of the dynamics in Eqn. 23. There are several different learning rules which can accomplish this. Mainly, we consider the Pseudo-inverse rule^39^ with synaptic weight matrix given by

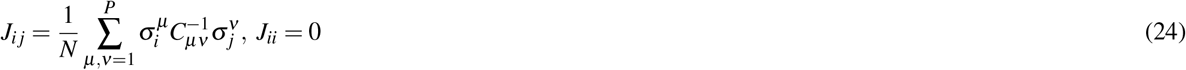

where the pattern overlap matrix *C*_*μν*_ is defined as

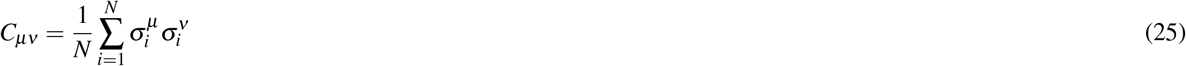

We also consider the Hebb rule given by

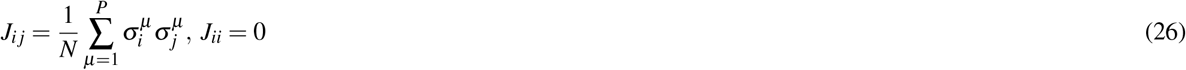

and the Storkey rule^**?**,63,64^ with *J*_*i j*_ given by

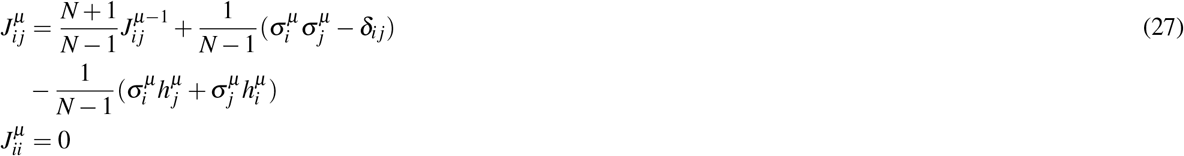

where,

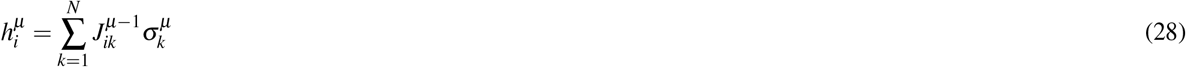

These learning rules differ in their capacity and the average size of the basins of attraction for memories at a given memory load *α*, further discussed Section 3.4 of the Supplementary Material.

#### Simulations

We simulate the memory storage, retrieval, and decoding processes by creating dictionaries of objects and attributes 𝒟^*a*^ and 𝒟^*b*^ where *a*’s and *b*’ s are random vectors with components drawn iid from a Gaussian distribution i.e. 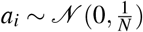. The size of these dictionaries is fixed to *D* = *L*_*max*_*N*, where *L*_*max*_ is the size of the largest structure being considered. In Fig. 1b we show the decoding error for several values of *L*_*max*_ and for Figs. 1c and 3 we set *L*_*max*_ = 30. We then use a subset of the dictionaries to create *P* knowledge structures with vector representations given by HRR encoding. These structures are then point-wise binarized and used to compute the synaptic weight matrix using the Pseudo-inverse rule unless otherwise stated. Since the encoding of the structures induces a similarity with individual relations *g*(*a, b*) rather than with *a* or *b* individually, we find that the same set of attributes {*b*_1_, *b*_2_, … *b*_*L*_} can be the same across several or all of the different knowledge structures while retaining the ability to decode the corresponding object 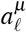 from retrieved structure 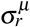. Hence, we consider the case in which the same attributes are used in each structure i.e.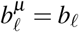.

We test the performance of the memory network by initializing the network in the state 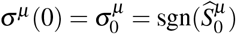 for each memory *μ* = 1, …, *P* where 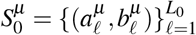 is the subset of *L*_0_ relations used to create a retrieval cue. We then evolve the network for *T* parallel updates, denoting the attractor reached by the network as 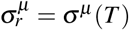, i.e., the retrieved state starting from partial cue of the *μ*-th structure. We define *m*^*μ*^ as the overlap between *σ*_*μ*_ and 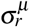 i.e.

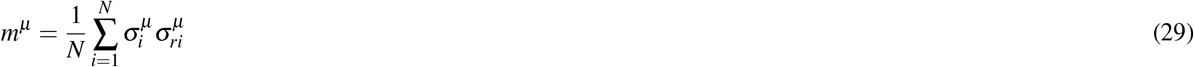

For each retrieved structure 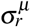, we use 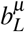, corresponding to a relation *not* contained in the initializing structure, to obtain an estimate 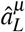 for 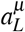, which then identified with the Dictionary element with which it has the highest overlap.

The Pseudo-inverse rule ensures that the basins of attraction for different structures are essentially identical regardless of potential differences in the overlap between different structures. In simulations, this allows us to consider each structure as an independent trial. The fraction of trials in which 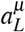 is incorrectly decoded from 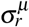 provides an empirical estimate of the decoding error *P*_*ε*_. We also construct an empirical SNR from Eqn. 20. Finally, we measure *m*^*μ*^ for each structure (Eqn. 29) to obtain an empirical distribution *p*(*m*) where the overlaps *m* are calculated for each memory in a trial and accumulated over many trials. The distribution *p*(*m*) does not appear to change if measured over multiple trials with different patterns or for multiple patterns within the same trial, which further supports the ability to consider each structure as a separate trial. The distribution *p*(*m*) is a statistical measure of the retrieval quality for structures of fixed size *L*, memory load *α*, and retrieval cue length *L*_0_.

#### Determination of *l*_*c*_

To determine *l*_*c*_ as a function of the various network parameters, we calculate the relation between *L*_0_*/L* and the average initial overlap *m*_0_ with the desired structure in the limit of large *N*, yielding

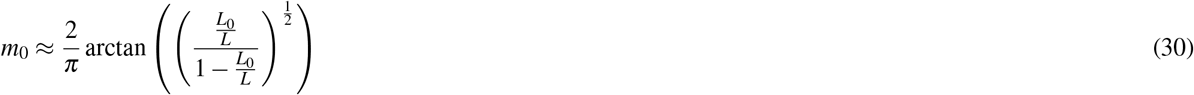

Further details of the derivation are provided in Supplementary Section 1.4 of the Supplementary Material. Using Eqn. 30, we write *l*_*c*_(*α*) in terms of *m*_*min*_(*α*) defined in the main text as

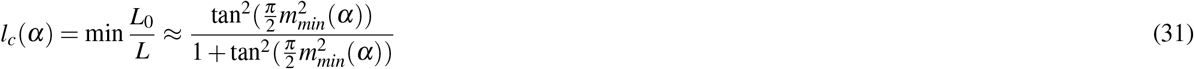

To determine *m*_*min*_(*α*) we resort to the Pseudo-inverse model with random binary patterns as memories^39^, which is simpler to simulate. Results relating *m*_*min*_(*α*) and *l*_*c*_ are shown in Supplementary Fig. 1.

#### Empirical Distribution of Overlaps

We find that the empirical distribution *p*(*m*) is bimodal and takes the general form

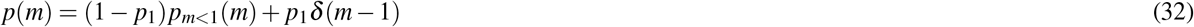

where *p*_1_ is the probability that a structure is perfectly retrieved from memory and *p*_*m<*1_(*m*) corresponds to the distribution of *m* for imperfectly retrieved memories.

The peak at *m* = 1 corresponds to trajectories converging to the target memories. This can be nonzero even when initial overlap *m*_0_ is outside the mean basin radius, indicating non-spherical basin shape. The second mode, peaked at 0 < *m* < 1 results from trajectories that converged to a fixed point outside the basin with a significant residual overlap with the memory. We characterize the shape of the distribution by the probability of *m* = 1, *p*_1_, the width of the lower *m* mode, *σ*_*m*_ and the mean of that mode, *m*\*. Results are shown in Supplementary Fig. 4a, for several values of *N* and two values of *α*.

As noted in^39,40^, the shape of the distribution *p*(*m*) is sensitive to finite size effects. To analyze these effects, we calculate *p*(*m*) for different sizes in a standard Pseudo-inverse model where the initial overlap *m*_0_ can essentially be varied continuously. For *m*_0_ > *m*_*min*_(*α*) almost all trials converge to the memorized pattern. For a range of values *m*_0_ < *m*_*min*_(*α*), *p*(*m*) is bimodal. We find that *p*(*m*) obtained from networks storing random patterns is very similar to the distribution obtain from networks storing structure memories, when the *m*_0_ and *L*_0_*/L* are related as in Eqn. 30. We find that for large *N, p*_1_ approaches a step function changing from zero to one as *m*_0_ crosses *m*_*min*_(*α*) = 1 − *R*(*α*). Near this transition, *p*_1_ can be approximated as

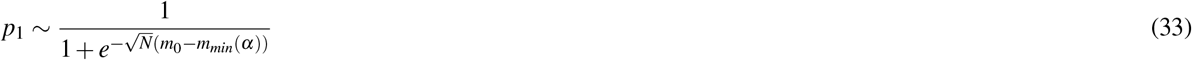

indicating that it converges to a step function exponentially fast with 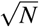. In addition, *σ*_*m*_ is very small outside the narrow transition regime of *m*_0_ and shrinks to zero everywhere as 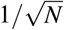. From this, we conclude that for *N* → ∞, *p*(*m*) becomes a *δ* function, which is either located at *m* = 1 for *m*_0_ > *m*_*min*_ or at a smaller value *m*\* which increases smoothly with *m*_0_, starting from zero and reaching 1 as *m*_0_ increases from zero to *m*_*min*_. Thus in large networks, the basins have a roughly spherical shape, such that virtually all initial conditions with *m*_0_ ≥ *m*_*min*_ converge to the memory, and all initial conditions with *m*_0_ < *m*_*min*_ converge to fixed points with partial overlap, *m*\*.

## Data availability

The data that support the findings of this study are available from the corresponding authors upon reasonable request.

## Acknowledgements

We thank Tankut Can, Naoki Hiratani, Mikhail Katkov, Kamesh Krishnamurthy, Andrew Saxe, Nimrod Shaham, and Misha Tsodyks for fruitful discussions. J.S. acknowledges support from the NSF through the Center for the Physics of Biological Function (PHY-1734030) and computational resources from the Princeton Research Computing at Princeton University, a consortium of groups led by the Princeton Institute for Computational Science and Engineering (PICSciE) and Office of Information Technology’s Research Computing. H.S. acknowledges support from the Swartz Program in Theoretical Neuroscience at Harvard, the Gatsby Charitable Foundation, and NIH grant NINDS (1U19NS104653).

## Author contributions statement

J.S. and H.S. designed the research, performed the research, wrote the main manuscript, and reviewed the full manuscript. J.S. prepared all figures and wrote the Supplementary Material.

## Additional information

The authors declare no competing interests.

## Supplementary Material for

### 1 Holographic Reduced Representation

#### 1.1 Overview

In holographic reduced representation (HRR)^1^, the binding operation *g* is circular convolution

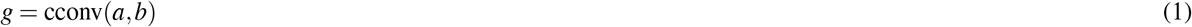

where the individual components of *g* are given by

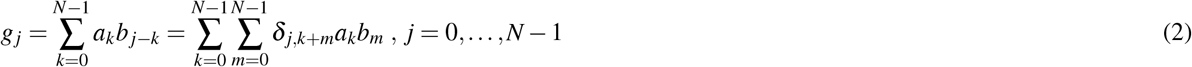

with all subscripts are defined modulo *N*. For objects and attributes *a, b* ∈ ℝ^*N*^, this creates a representation *g*(*a, b*) ∈ ℝ^*N*^. The circular convolution operation is both associative and commutative, i.e. cconv(*a, b*) = cconv(*b, a*).

The corresponding decoding operation is given by the circular correlation

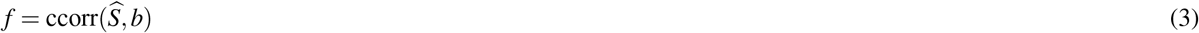

where the individual components of *f* are given by

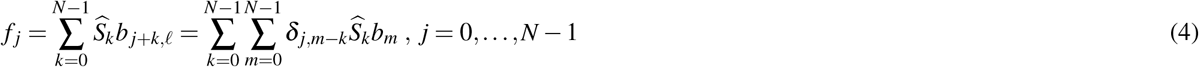

In general the circular correlation operation is neither associative nor commutative. However, it can be related to circular convolution by defining the involution of a vector *a* as *a*\* where 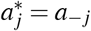. Then ccorr(*a, b*) = cconv(*a*\*, *b*) and *a*\* as serves as an approximate inverse to *a* when both *a* and *b* are random vectors.

#### 1.2 Clean-up and SNR

To quantify the effect of the ML clean-up operation described in the main text, we compute various statistics of the estimator *â*_𝓁_ for the corresponding object *a*_𝓁_ decoded from a relational structure 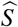 using the HRR unbinding operation. We consider the case where all objects and attributes are random vectors with components drawn iid, i.e., 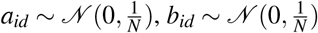, and decoding is done from a dictionary of size *D*.

Since the encoding operation is permutation invariant, the statistics for all *â*_𝓁_ decoded from the full structure should be the same. For simplicity we set 𝓁 = 1 and consider *â*_1_. We index dictionary items so that indices 𝓁 < *L* correspond to the dictionary items contained in the structure 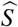 and indices *L* < 𝓁 < *D* corresponds to dictionary items that are not contained in 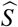. Note that

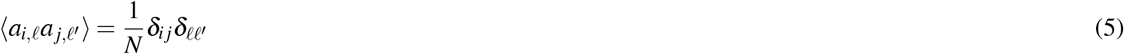

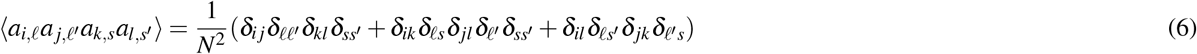

Using the expression for the unbinding operation in Eqn. 4, we can express the estimator *â*_1_ as

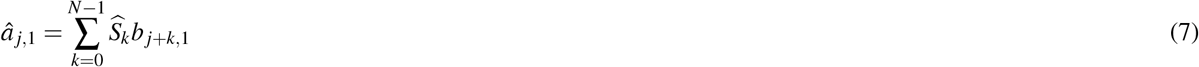

Using Eqn. 2, we can express this in terms of *a*_𝓁_ ‘s and *a*_𝓁_ ‘s, and *â*_*j*,1_ as

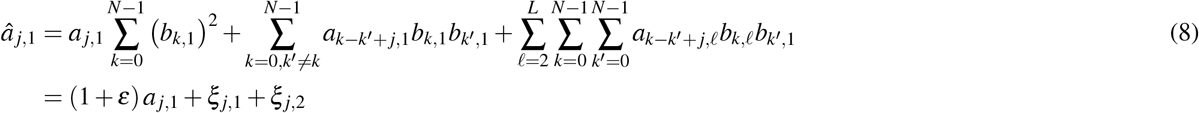

where we have defined the three noise terms as

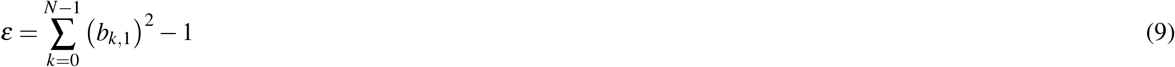

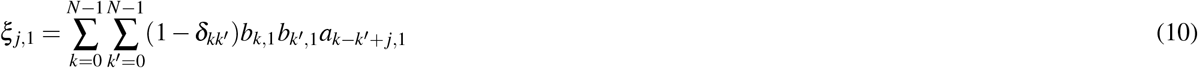

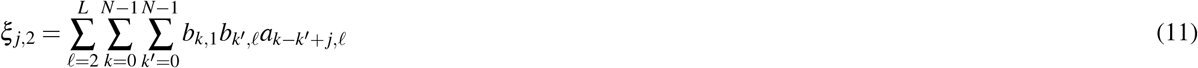

Here, *ε* is noise due to deviations in the normalization of *a, ξ*_*j*,1_ is noise from interference elements of different elements of the cue within the same attribute *b*_1_, and *ξ*_*j*,2_ and *ξ*_*j*,2_ are noise coming from interference between attribute *b*_1_ with all other attributes *b* contained in the structure. Since components of *a*_1_ appear an odd number of times in each term of the sums in *ξ*_*j*,1_ and *ξ*_*j*,2_, we conclude that ⟨ *ξ*_*j*,1_ ⟩ = ⟨ *ξ*_*j*,2_ ⟩ = 0. Likewise, ⟨ *ε* ⟩ = 0. Using Eqns. 5 and 6, we find that the second moments and correlation of the two noise terms are given by

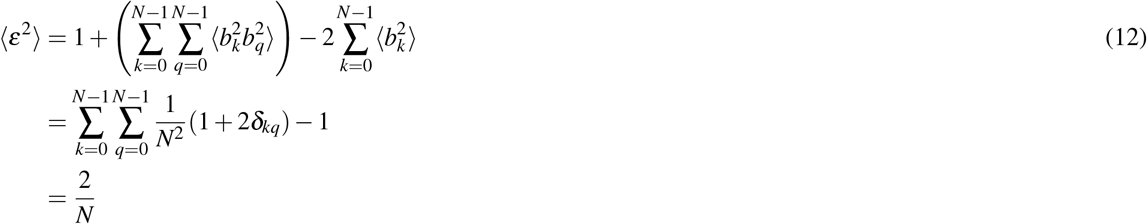

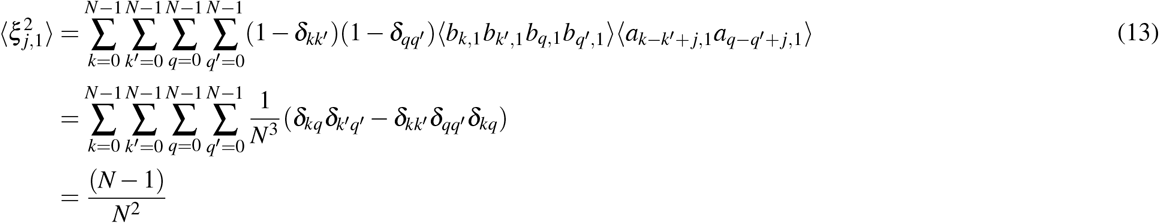

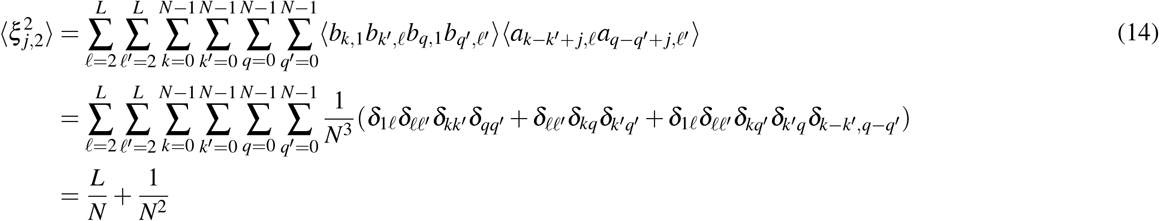

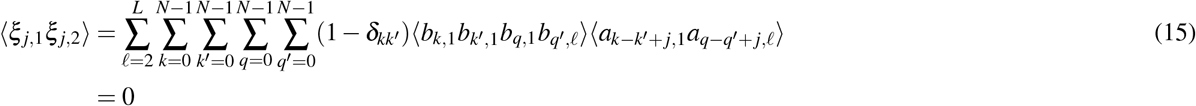

and from Eqn. 8, we see the noise in each component of the estimator is

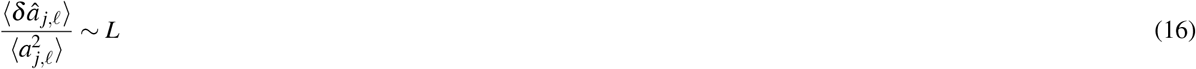

We now calculate the first two moments of the overlap of the estimator with the correct pattern *a*_*d*_ · *â*_1_ to obtain the SNR for HRR defined in Eqn. 20 of Methods. *a*_*d*_ · *â*_1_ can be expressed as

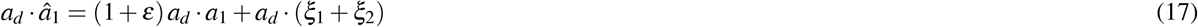

We see from Eqn. 17 that ⟨ *a*_1_ · *â*_1_ ⟩ = 1 and ⟨ *a*_*d*_ · *â*_1_ ⟩ = 0, so the estimator is unbiased. The second moment is given by

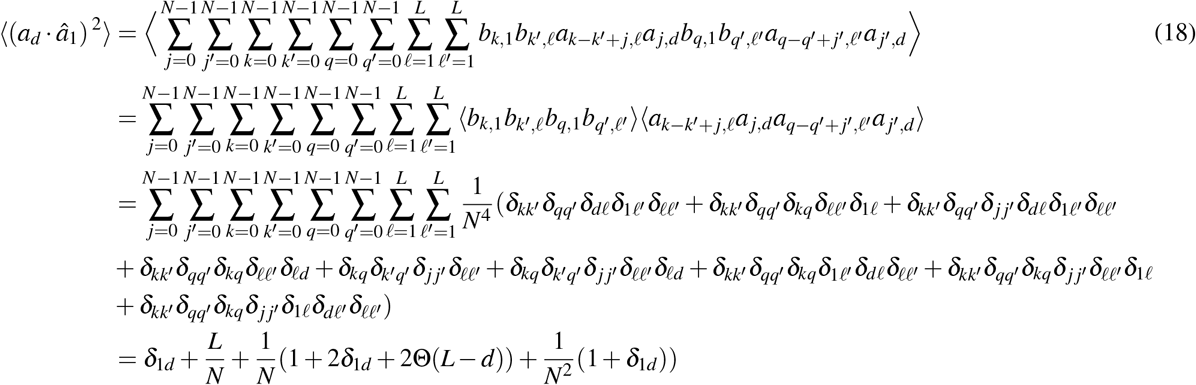

To summarize

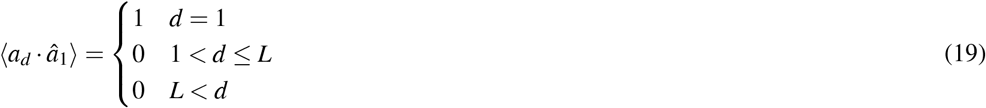

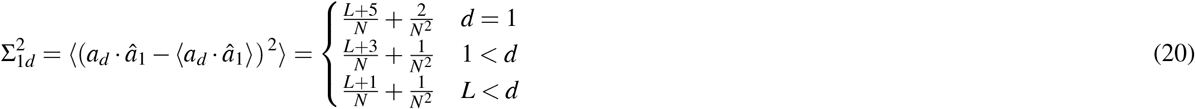

For *L* ≪ *N*, the SNR for overlaps defined in Eqn. 20 of the main text is then approximately

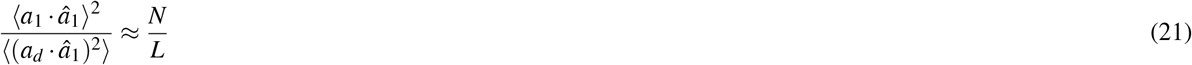

#### 1.3 Binarization and Empirical SNR

When decoding from a binarized structure 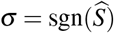, the estimator *â*_1_ is given by

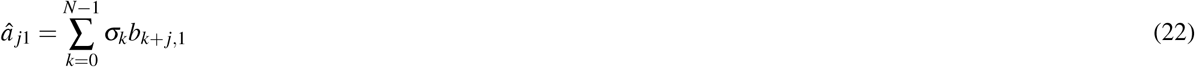

where

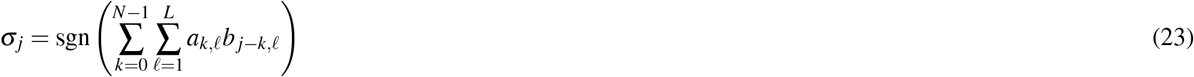

To compute the SNR, we start by evaluating the overlap between the estimator *â*_1_ with items in the dictionary ⟨ *a*_*d*_ · *â*_1_ ⟩ which can be expressed as

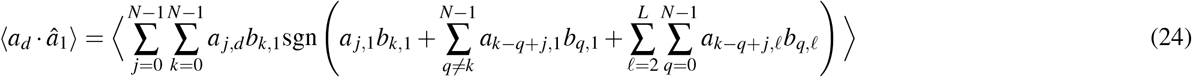

While individual products of elements *a*_𝓁,*i*_*b*_𝓁, *j*_ are not Gaussian distributed, the expectation ⟨ *a*_*d*_ · *â*_1_ ⟩ is the sum of *N*^2^ random variables with correlations only occurring at higher order as the expectation of the product of different elements of each circular convolution is zero. Since all terms contained in the sums over indiced *j* and *k* in Eqn. 24 are only correlated at higher order, we can approximate each term in the sum as independent which gives us

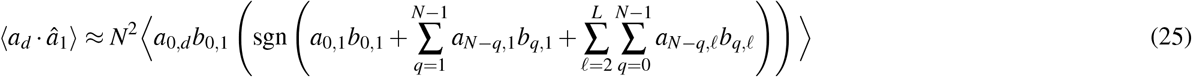

We now define the following three random variables

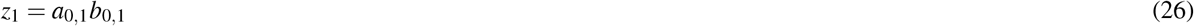

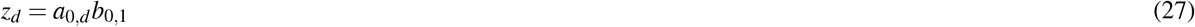

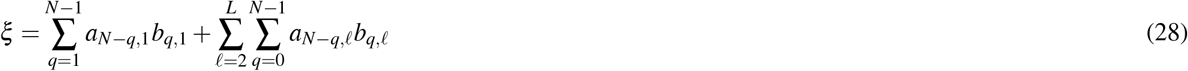

where *x*_1_ is a signal term and *ξ* is a noise term within the sign function. ⟨*z*_1_ ⟩= ⟨*z*_2_ ⟩= ⟨*ξ* ⟩= 0 and the variances are given by

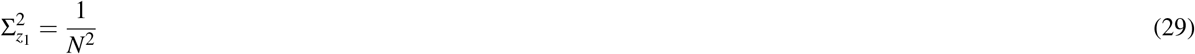

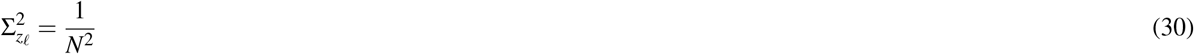

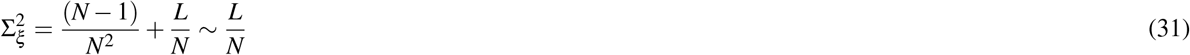

For *d* = 1, *z*_1_ = *z*_𝓁_ = *z* and *L* ≫ 1 we can treat *z* and *ξ* as Gaussian. In this approximation ⟨ *a*_*d*_ · *â*_1_ ⟩ becomes

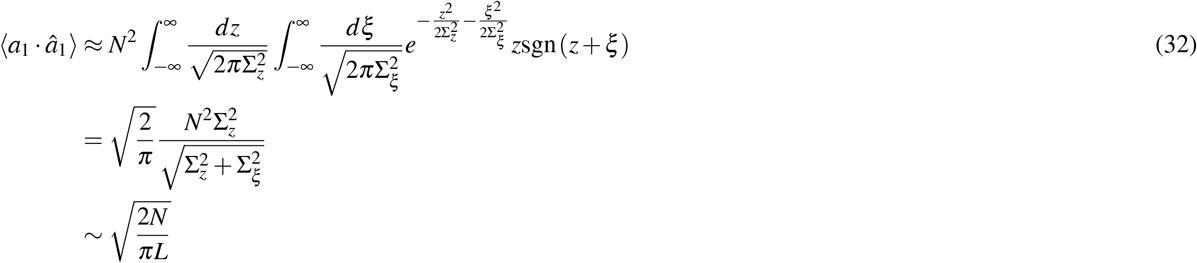

Likewise, the second moment is given by

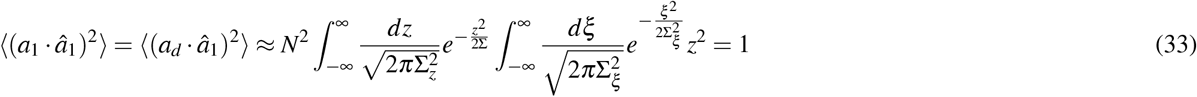

Putting Eqns. 32 and 33 together, the SNR of overlap when decoding *a*_1_ from the binarized structure *σ* is given by

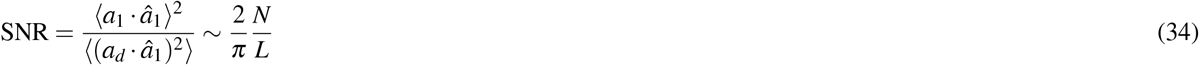

We see from Eqn. 21 that binarizing 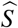 has decreased the SNR by a constant factor of 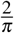.

#### 1.4 Memory Initialization

We now determine the average initial overlap *m*_0_ between a binarized retrieval structure *σ*_0_ containing *L*_0_ of the *L* relations in the binarized full structure *σ* in the case of random *a*_𝓁_ ‘s and *b*_𝓁_ ‘s are random vectors with components drawn iid as before. An expression for the average overlap *m*_0_ between *σ*_0_ and *σ* can be written as

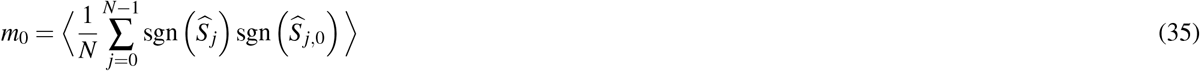

The overlap between components of the unbinarized structures 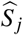 and 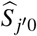 with HRR binding is given by

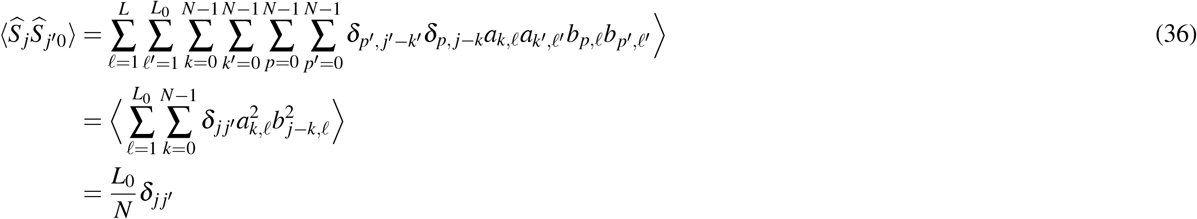

For large *N* we can approximate the sum in Eqn. 35 by treating the individual terms as as independent since there are no lower order correlations between terms. We can then define the random variables 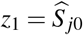 and 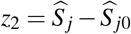 as approximately Gaussian distributed with zero mean and variances 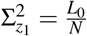 and 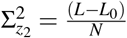 respectively. Then, Eqn. 35 can be approximated as

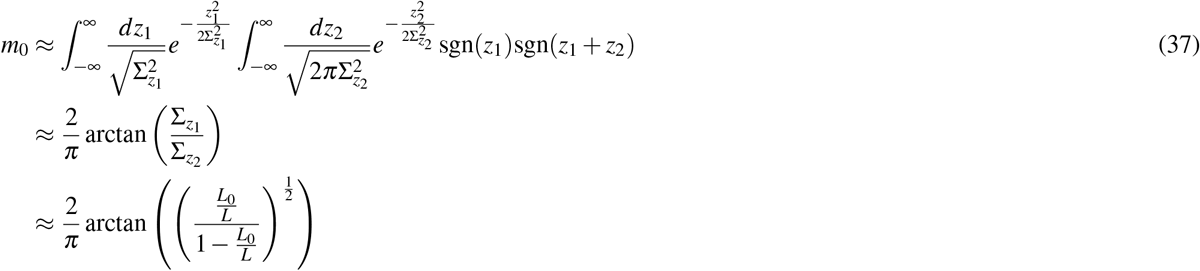

For *L*_0_ ≪ *L* Eqn. 37 can be further approximated as

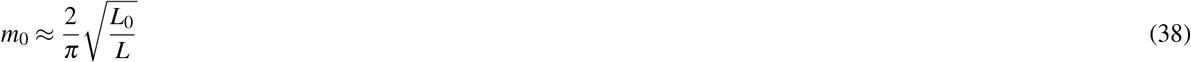

This implies that creating retrieval structures with *L*_0_ out of *L* relations will result in retrieval cues with average overlap *m*_0_ with the full memorized structure. The variance of the distribution of initial overlaps will vanish as *N* → ∞.

We use this relation to determine *l*_*c*_(*α*) as a function of *L*_0_*/L* using the value of *m*_*min*_(*α*) obtained for a Pseudo-inverse model with random binary patterns as memories^2^. Results for *m*_*min*_(*α*) are shown in Fig. 1a demonstrating a linear increase in the range *α* < 0.5. Substituting these results in Eqn. 31 closely predicts the value of *l*_*c*_(*α*) as shown in Fig. 1b.

**Figure 1.**
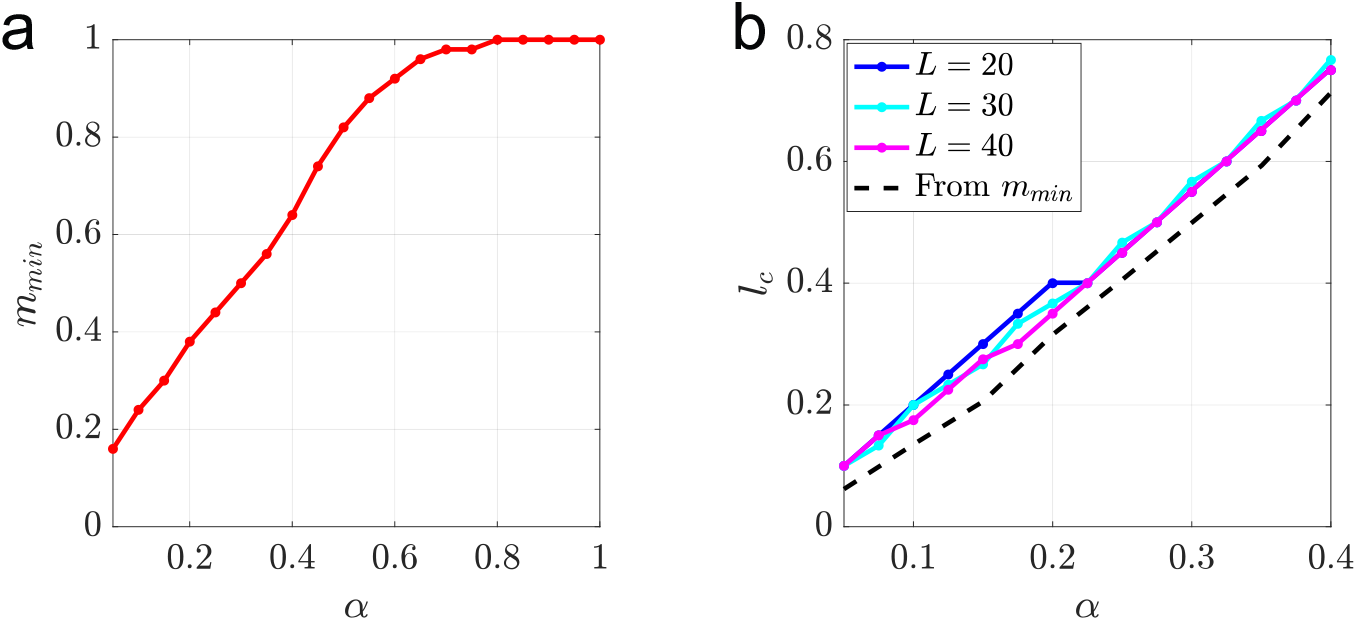
(a) *m*_*min*_ shown as a function of *α* for a Pseudo-inverse network storing with random patterns. (b) For structured memories, *m*_*min*_ corresponds to a minimum fraction of relations needed obtain a final overlap *m* = 1, shown for *L* = 20, 30, 40 along with the value predicted by *m*_*min*_ from the expression in Eqn. 31. *N* = 1000 and *T* = 20 parallel updates are used in memory retrieval for both figures and the averages are computed over 50 trials.

### 2 Decoding Error

We derive a good approximation for the ML decoding error in terms of *D*, the size of the decoding dictionary, and SNR, the signal-to-noise ratio of overlaps defined in Eqn. 20 of Methods. The overlap 𝒪_*d*1_ between a dictionary item *a*_*d*_ with the estimator *â*_1_ for *a*_1_ decoded from a structure 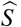 is given by

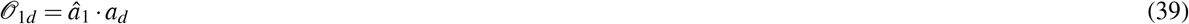

where as in the previous section, we index items so that *d* ≤ *L* corresponds to overlap with patterns within 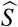 and *L* < *d* corresponds to overlaps with other items in the Dictionary not contained in 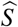. For ML decoding, an error occurs if max(𝒪_12_, …, 𝒪_1*D*_) ≥ 𝒪_11_. The probability of error *P*_*ε*_ is then given by

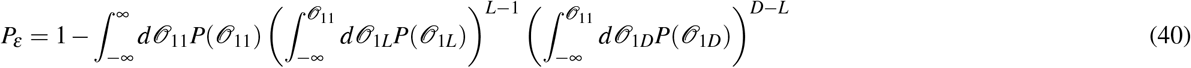

For large *N*, each 𝒪_1*d*_ essentially behaves as an independent Gaussian random variable with mean *μ*_1*d*_ and variance Σ_1*d*_. For unbiased decoding schemes, *μ*_1*L*_ = *μ*_1*D*_ = 0. In general, we can consider decoding from a structure containing errors by setting *μ*_11_ = *μ*, where *μ* is related to the overlap of the corrupted structure with the correct structure. Then *P*_*ε*_ can be very well approximated as

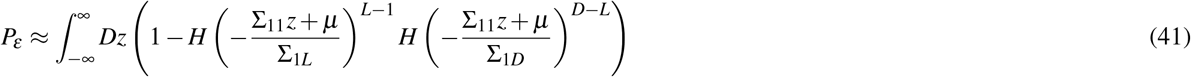

where 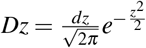.

For HRR Σ_11_ ≈ Σ_1*L*_ ≈ Σ_1*D*_ to leading order in 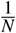. Defining Σ to be the leading order term in the variance, we obtain

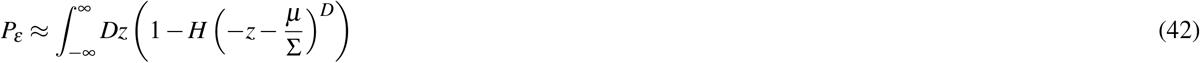

Identifying 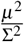 with the SNR of overlaps, Eqn. 42 becomes

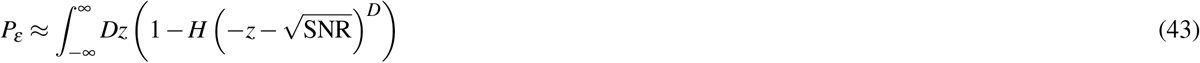

given in the main text.

#### 2.1 Limitations for Good Decoding

We can understand the dependence of *P*_*ε*_ on the SNR and D in the large SNR regime by obtaining a saddlepoint approximation for the expression for *P*_*ε*_ given in Eqn. 43. We start by rewriting Eqn. 43 as

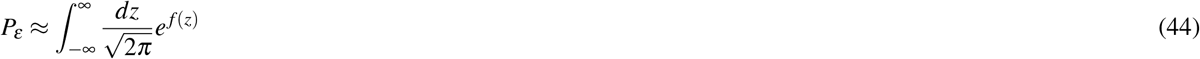

where *f* (*z*) is given by

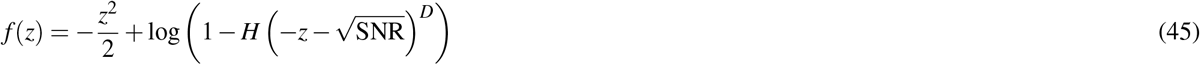

From Eqn. 42, we see *P*_*ε*_ ≪ 1 for 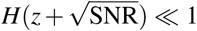 as 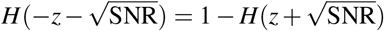. So we are interested in the regime where 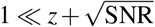. In this regime, *f* (*z*) is very large, so the integral in *P*_*ε*_ can be approximated its saddlepoint value

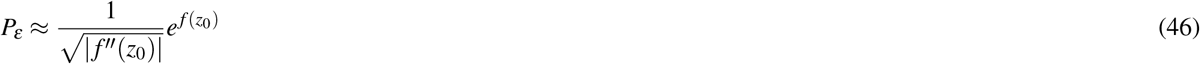

where *z*_0_ is given by the solution of

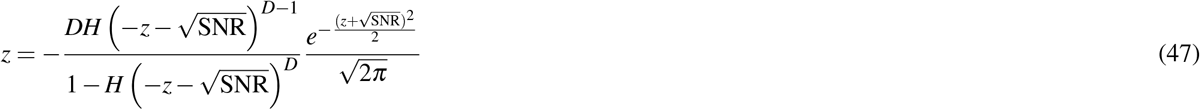

In this regime we can make the approximation

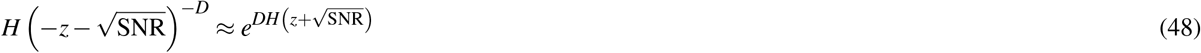

which allows us to approximate Eqn. 47 as

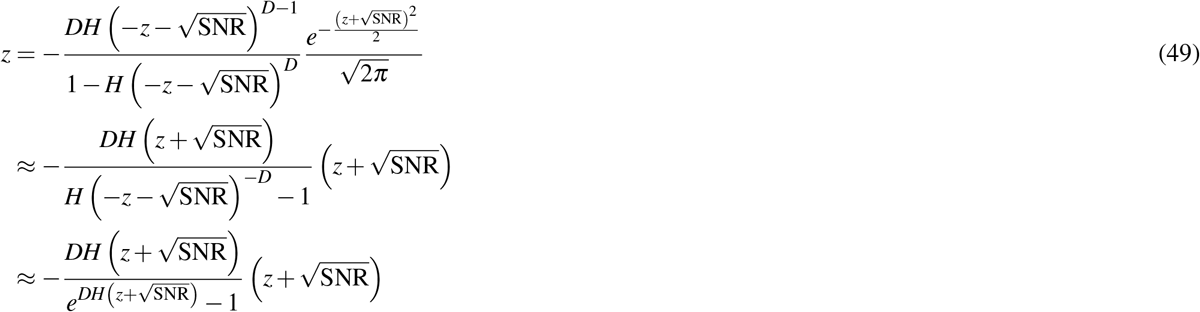

where for 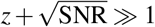 we used

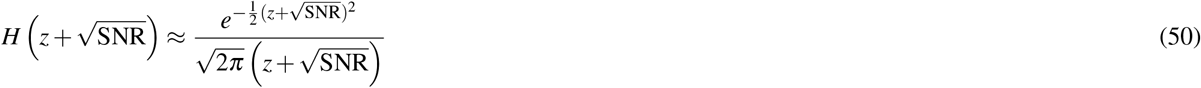

From Eqn. 49 we see that for 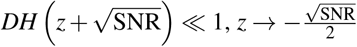 and 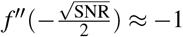. The saddlepoint approximation for the error is then

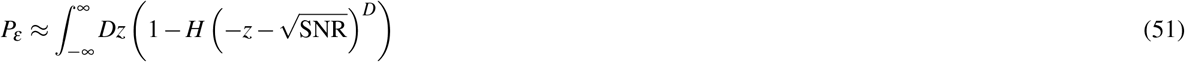

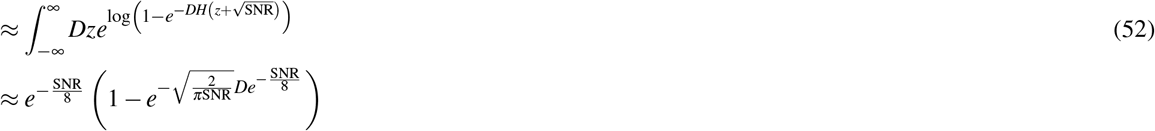

This can be further approximated as

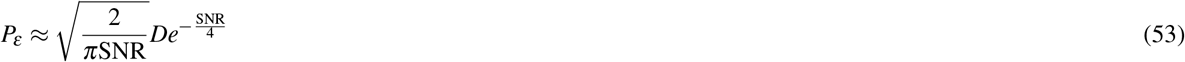

We can insert the SNR of overlaps into Eqn. 53 to determine the limits on the size of *D* for good decoding. For HRR the approximation for the error gives us

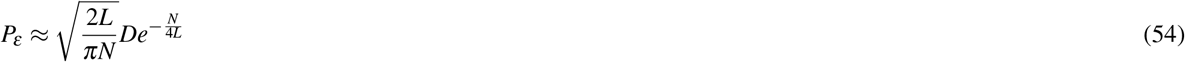

which implies that for an error of *δ* ≪ 1

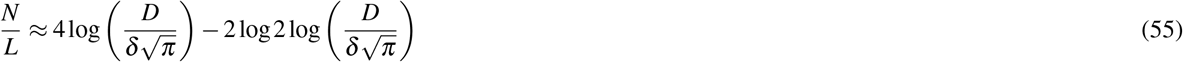

From this, we expect the error to be small as long and SNR is large and *D* is polynomial, and not exponential, in *N*.

#### 2.2 Decoding After Memory Retrieval

To analyze the effect of memory retrieval on the decoding error, we start by considering decoding from a degraded binarized structure which is a Hamming distance 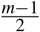 from the uncorrupted binarized structure. In this case, the effective SNR is found by making the replacement ⟨ *a*_*d*_ · *â*_1_ ⟩ in Eqn. 34. From this, we see that SNR(*m*) should take the form

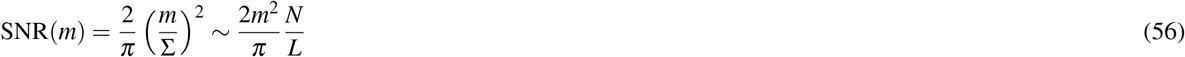

When *â*_1_ is decoded from an imperfectly retrieved structured from memory, we can instead make the replacement 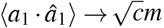. Here, *c* ∼ *O*(1) is a constant factor accounting for differences in overlap of the structure with relations within and outside the retrieval cue. Then SNR(*m*) takes the modified form

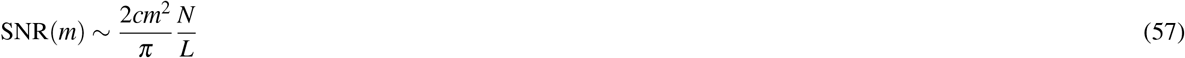

The decoding error after retrieval from memory is then given by

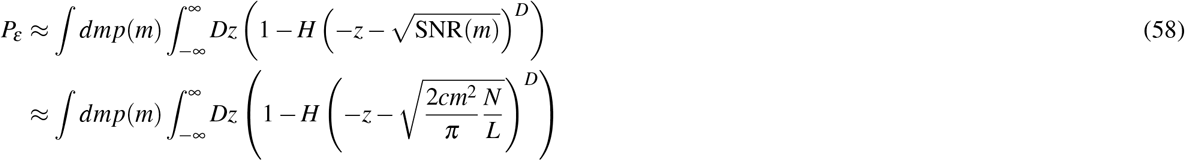

where *p*(*m*) takes the same form as in Methods. For *N* → ∞, *p*(*m*) → *δ* (*m* − *m*\*) and the decoding error approaches

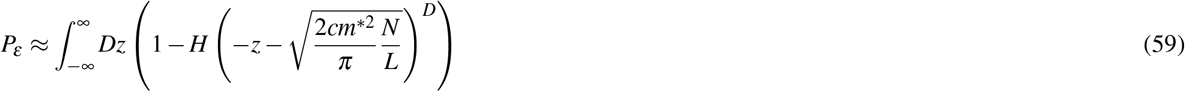

### 3 Insights from Random Patterns

#### 3.1 Network Order Parameters

For generic Hopfield networks, we can characterize the quality of memory retrieval by formally defining three network order parameters which quantify the overlap of the network state with the stored memories at each time step. The first is

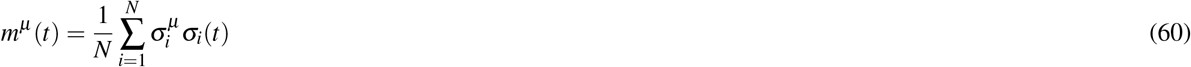

which represents the overlap of the network state *σ* (*t*) with the corresponding pattern *σ*^*μ*^ at time *t*. If the initial state of the network has an *O*(1) with a small number of patterns, i.e., *m*_0_ = *m*^*μ*^ (0), the memory retrieval process can be sufficiently described by including an additional order parameter

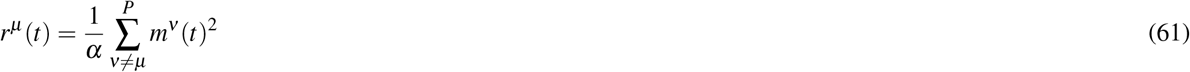

which represents the overlap of the state with all patterns except for *μ*. We can also define

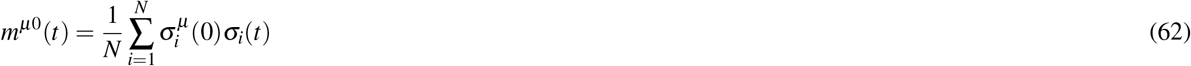

as the overlap of the state at time *t* with the initial state of the network after attempting to retrieve *σ*^*μ*^ where the overlap between *σ*^*μ*^ (0) and *σ*^*μ*^ is *m*_0_. While *m*^0^ is typically not considered for random patterns, it becomes relevant in the case of structured knowledge, where the retrieval cue can be constructed from a subset of relations rather than simply a corrupted version of a memory.

For large *N*, the distributions of order parameters over fixed retrieval conditions, i.e., *p*(*m*), *p*(*r*) and *p*(*m*^0^), are sharply peaked at the average values, denoted by ⟨*m* ⟩, ⟨*r* ⟩, and ⟨*m*^0^ ⟩. In the limit *N* → ∞, these quantities depend solely on *m*_0_ and *α*.

#### 3.2 Overlap Distributions

As discussed in Methods, the empirical distribution *p*(*m*) is bimodal and takes the general form

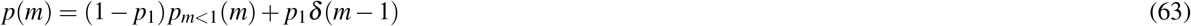

where *p*_1_ is the probability that a structure is perfectly retrieved from memory and *p*_*m<*1_(*m*) corresponds to the distribution of *m* for imperfectly retrieved memories. The shape of the distribution is characterized by *p*_1_, the width of the lower *m* mode, *σ*_*m*_ and the mean of that mode, *m*\*. These quantities all depend on the initial overlap *m*_0_ used to retrieve the memory Results for *p*_1_, *m*\* and *σ*_*m*_ are shown in Fig. 2a, for several values of *N* and two values of *α*.

We also compare the overlap distributions for structures of length *L* with retrieval cue of length *L*_0_ = 2 with the overlap distribution for a Pseudo-inverse network with random patterns for the corresponding values of *m*_0_ given in Eqn. 35 in Fig. 3.

#### 3.3 Retrieval Dynamics

When retrieving structures from memory (in the absence of noise), the update equation for the state of each neuron at time *t* is given by

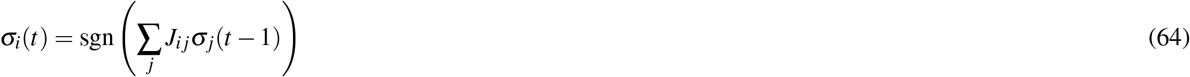

For all of the simulations in the main text, we consider parallel updates where all of the neurons 1 ≤ *i* ≤ *N*. We find that serial updates give qualitatively similar results to parallel updates. In general, for large *N* we find

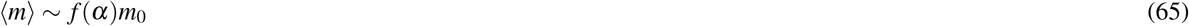

where ⟨*m* ⟩ → *m*\* as *N* → ∞. We find that serial updates obey the form in Eqn. 65 with a slightly smaller value of the coefficient *f* (*α*). Additionally, for serial dynamics, we can consider the robustness of our results under addition of noise in the updates. To do this, we use the Metropolis algorithm to obtain the final equilibrium state, where the amplitude of the noise is controlled by the inverse temperature *β*. At each update, the acceptance probability is given by

**Figure 2.**
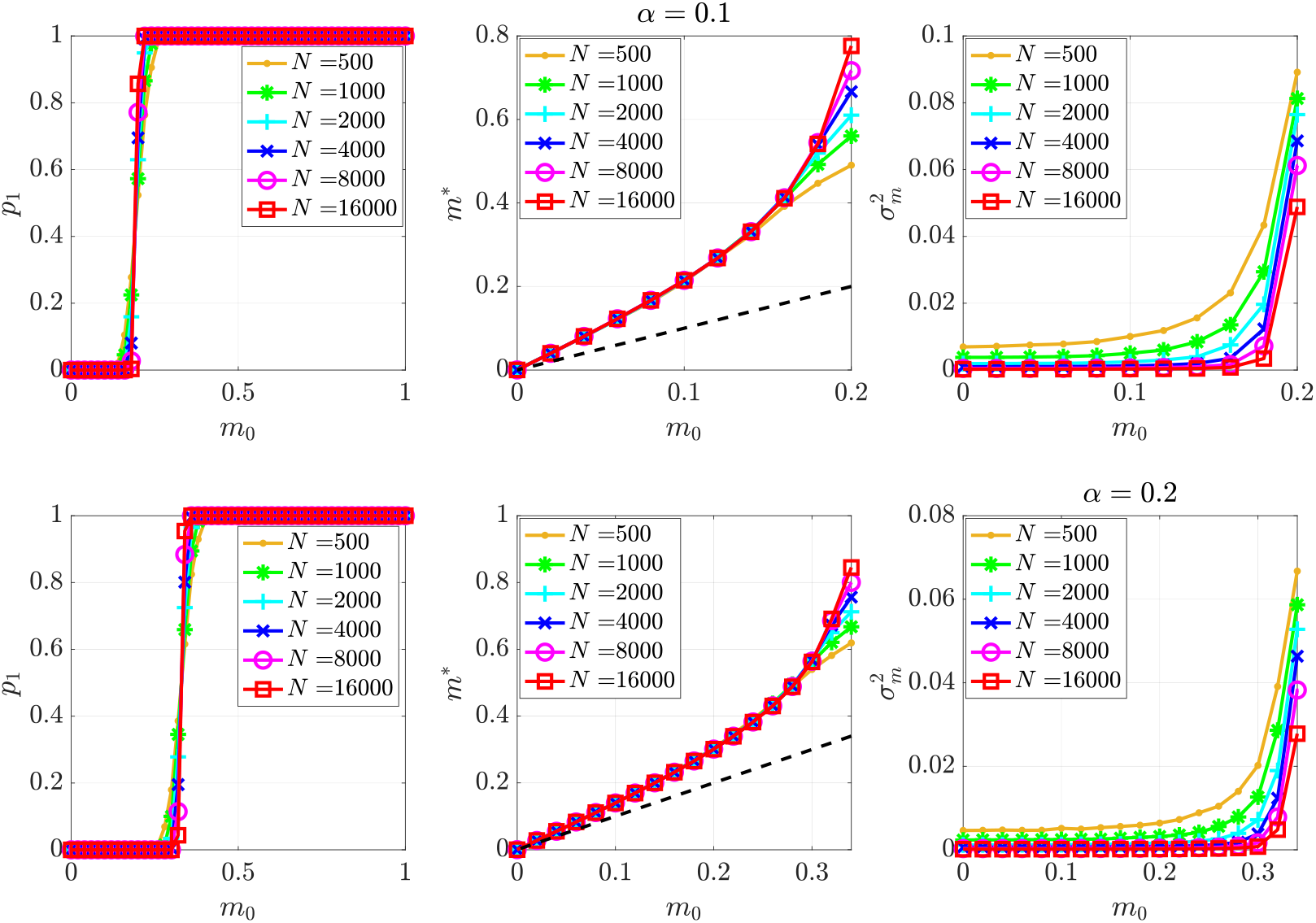
*p*_1_, *m*\*, and 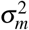 are shown as functions of the initial overlap *m*_0_ for several values of *N* for Pseudo-inverse networks storing random memories for two values of the memory load *α*. The dotted black line in the middle figures show that *m*\* > *m*_0_ for these memory loads even when the initial state is far outside of the basin of attraction. *T* = 20 parallel updates are used for memory retrieval and averages are performed over 50 trials.

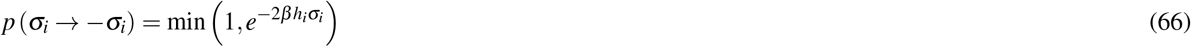

We see that Eqn. 66 becomes equivalent to Eqn. 64 for the noiseless case *β* = ∞. In Fig. 4, we show the average overlap *m* of the network state with the cued memory as a function of time for several different initial overlaps *m*_0_ retrieved through parallel dynamics and serial dynamics for several different values of *β*. In Fig. 5 we show ⟨*m* ⟩, ⟨*r* ⟩, ⟨*m*^0^ ⟩ as function of retrieval cue overlap *m*_0_. We see that the highest overlap is achieved for parallel updates when the serial updates are done in random order (alternative orders are discussed in^2^). However, the results are qualitatively similar for the different dynamics shown, demonstrating that small amounts of noise have little effect.

In Fig. 5a and c, we see that outside of the basin of attraction of the cued memory, i.e., values of *m*_0_ where ⟨*m* ⟩ < 1, the final network state retains some memory of the initial state reflected by ⟨*m*^0^ ⟩ > *m*_0_.

#### 3.4 Comparing Learning Rules

In the main text, we considered storing structured memories in recurrent neural networks where the synaptic weights are set via the Pseudo-inverse rule. This learning rule fully de-correlates linearly independent memories so that each memory is a perfect fixed point for *α* < 1. However, the Pseudo-inverse rule is both non-local and non-incremental. This makes it unlikely to be biologically implemented in a straightforward manner.

We now consider two learning rules that are both local and incremental. The first is the Hebb rule and the second is the Storkey rule proposed in^**?**,3,4^ (see Eqns. (26)–(28) in Methods). The Storkey rule can be viewed as a biologically plausible approximation to the Pseudo-inverse rule for small *α*^4^. As a result, the storage capacity of this rule *α*_*c*_ ≈ 0.4 is significantly higher than the Hebb rule (*α*_*c*_ ≈ 0.14) and the basins of attraction are larger and more even across different memories^**?**^.

The storage and retrieval of random memories in Hopfield networks near saturation, i.e., when the number of memories *P* scaling linearly in *N*, is limited by interference between different memories^5^. The overlaps between different memories are characterized by the following matrix

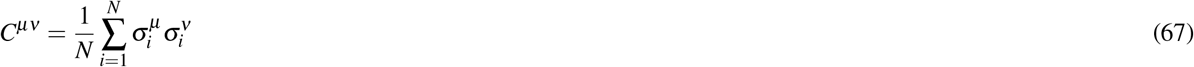

Note that *C*^*μν*^ is a *P* × *P* matrix that has the form of a sample covariance matrix.

**Figure 3.**
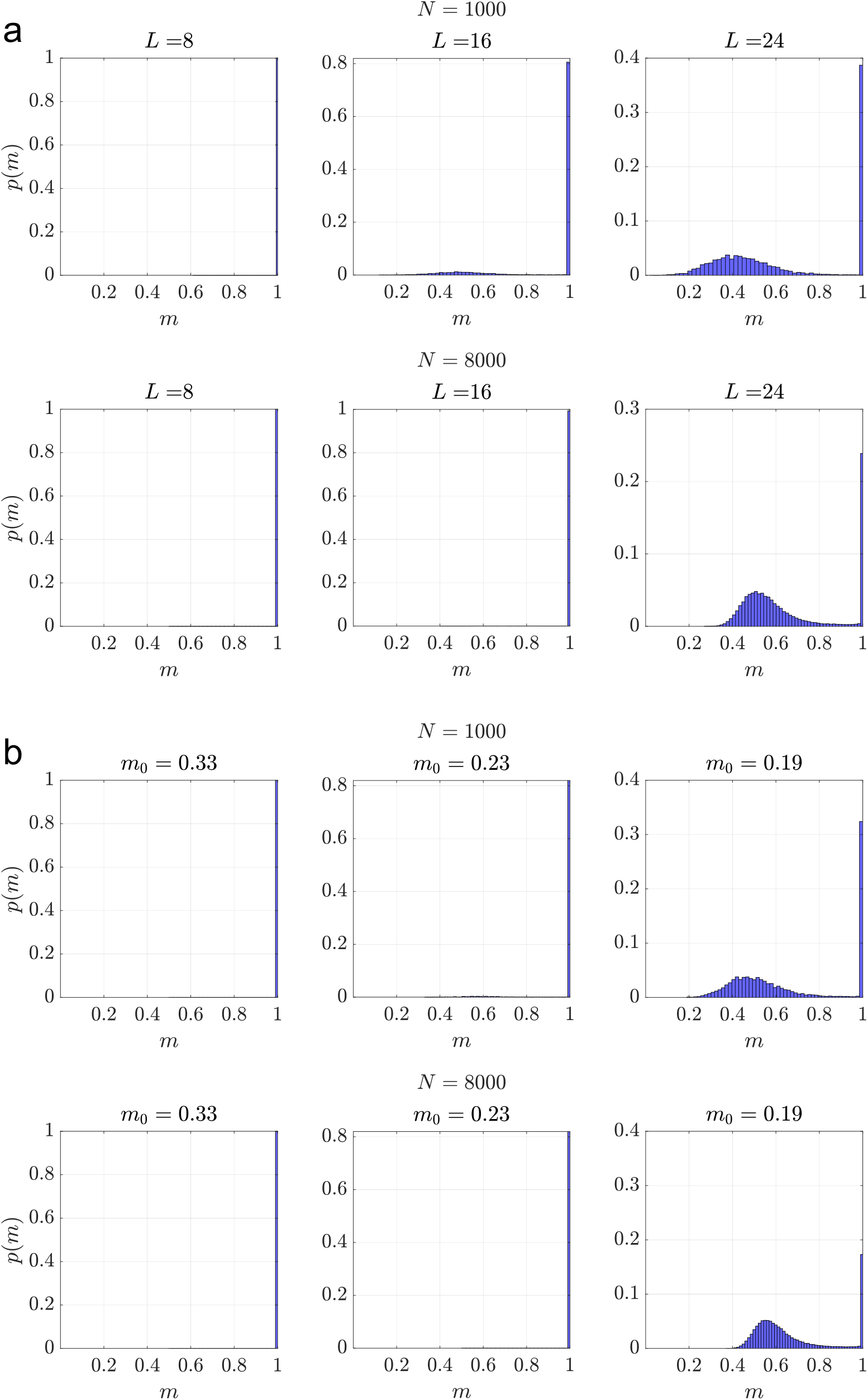
(a) Histograms of the final overlaps *m* for structures of various lengths of *L* for *α* = 0.1 and *L*_0_ = 2 and 100 trials. *T* = 20 parallel updates are used to retrieve memories. The top panel shows *N* = 1000 and bottom shows *N* = 8000. (b) Histograms for values of *m*_0_ obtained from Eqn. 37 corresponding to *L*_0_ = 2 and *L* = 8, 16, 24 (left to right).

**Figure 4.**
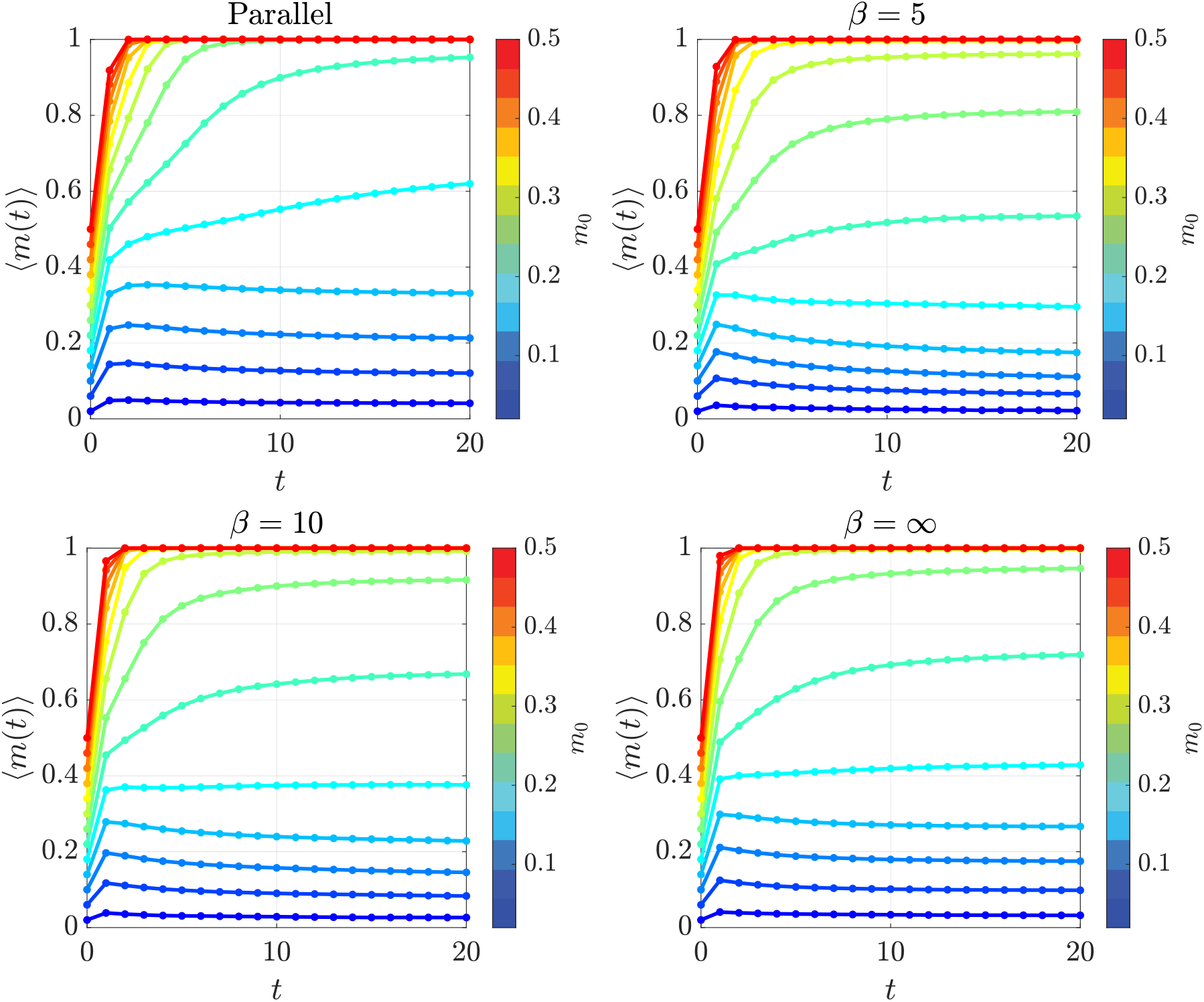
Evolution of the averaged overlap order parameter ⟨*m* ⟩ over *T* = 20 updates compared for parallel updates serial dynamics at zero noise (*β* = ∞), and serial dynamics with noise amplitude determined by inverse temperature *β* = 5, 10, for several different initial conditions *m*_0_. The Pseudo-inverse rule is used to store the memories. *α* = 0.1, *N* = 1000, and 50 trials are used for all figures.

**Figure 5.**
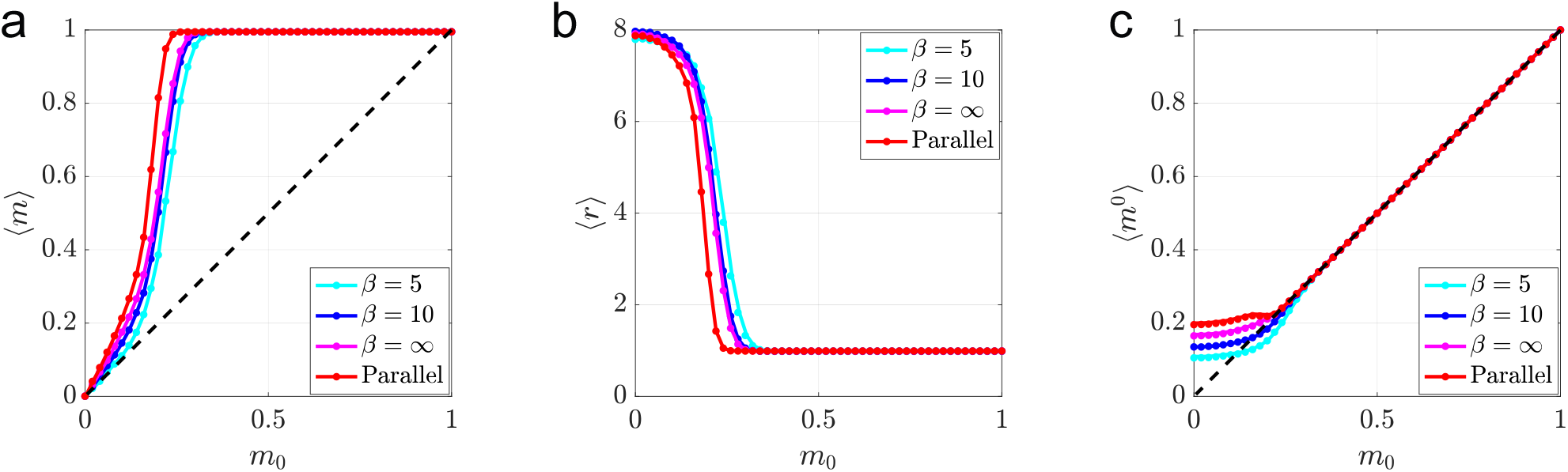
Comparison of averaged order parameters ⟨*m* ⟩ (a), ⟨*r* ⟩ (b), and ⟨*m*^0^ ⟩ (C), after *T* = 20 updates as functions of retrieval cue overlap *m*_0_ are compared for parallel updates and serial updates for various values of *β*. The dashed line in A and C is shown to compare the value of the final overlap with the cued memory *m* and the initial state *m*^0^ to the initial overlap with the cued memory. The Pseudo-inverse rule is used to store memories and *α* = 0.1, *N* = 1000, 50 trials are used for all figures.

We can compare the Pseudo-inverse, Hebb, and Storkey rules by looking at the forms of the local fields given by

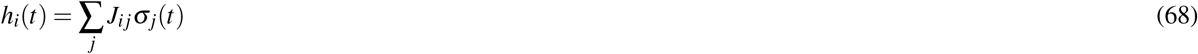

From Eqn. 64, we see that a memory is a fixed point if 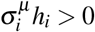 for all *i* = 1, …, *N*. The elimination of the self-coupling term *J*_*ii*_ greatly increases the basins of attraction of the pattern. In the limit *N, P* → ∞ this can be accomplished by replacing *J*_*i j*_ with *J*_*i j*_ − *αδ*_*i j*_^2^. The interference between memories is contained in the noise term in the local field.

We start by considering the Hebb rule, with synaptic weight matrix given by

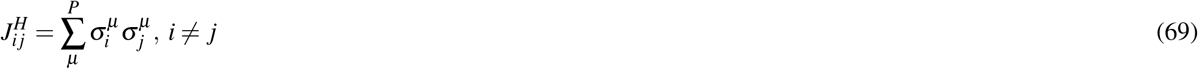

The local field for the Hebb rule can be expressed as

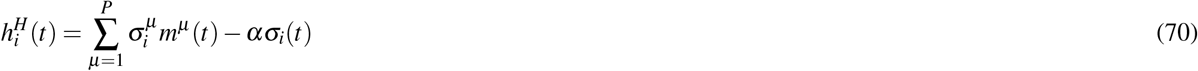

Next we consider the the Pseudo-inverse rule with synaptic weight matrix expressed in terms of *C*^*μν*^ as

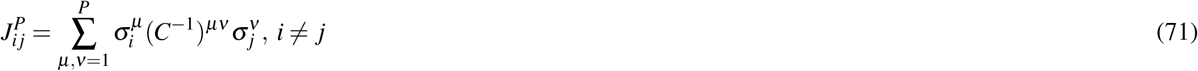

It is useful to decompose the state of the network *σ*_*i*_(*t*) in two parts as follows

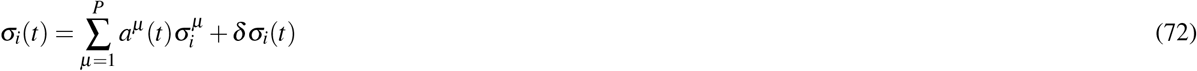

where *δσ*_*i*_(*t*) is orthogonal to all of the patterns 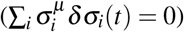 and *a*^*μ*^ is related to the order parameter *m*^*μ*^ (*t*) via

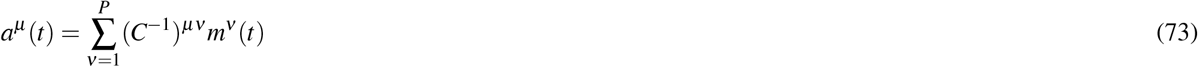

The local field for the Pseudo-inverse rule can then be expressed as

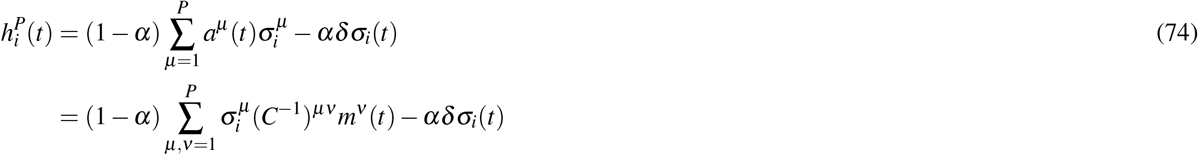

We can see how Eqn. 74 suppresses the effects of overlaps by considering the state 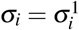. Then *a*^*μ*^ = *δ*_*μ*1_ regardless of any correlations of the memories and 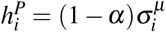. This implies that each memory is an eigenvector of 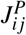 with eigenvalue (1 − *α*) so that all memories are perfect fixed points for *α* ≥ 1. By contrast, we see from Eqn. 70 that 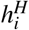 becomes

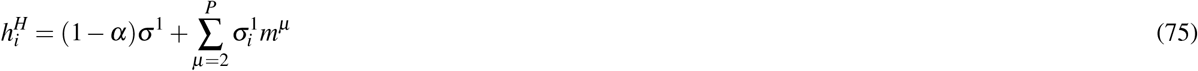

contains additional noise terms of 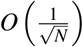 because of overlaps with the other patterns. This noise reduces the capacity of the network as well as the size of the basins of attraction for each memory which are both further reduced if the patterns are not random and contain correlations.

A simplified form of the Storkey rule, discussed in^**?**,4,6^, is given by

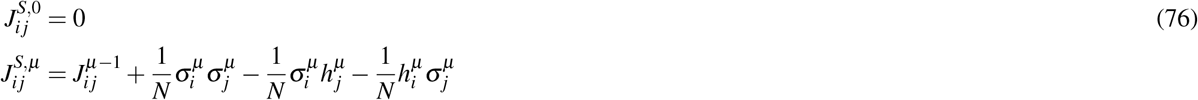

where

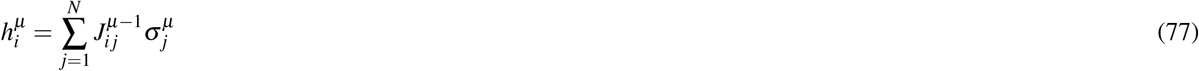

Following the analysis in^6^ We can find a more compact approximation for 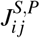. We start with *P* = 1. Then 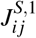 is given by

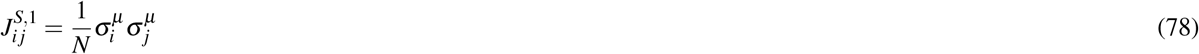

For *P* = 2 we have

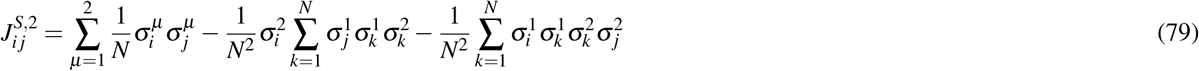

For *P* memories, keeping terms up to 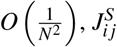 can be expressed as

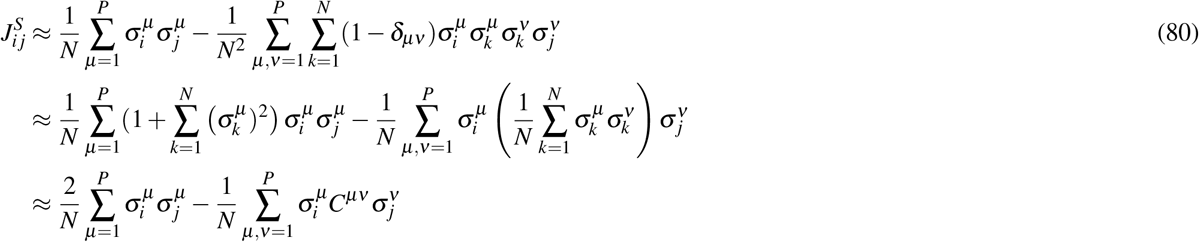

We can relate 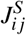 to the Pseudo-inverse rule given in Eqn. 71 by rewriting 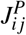 using the following series expansion of *C*^−1^

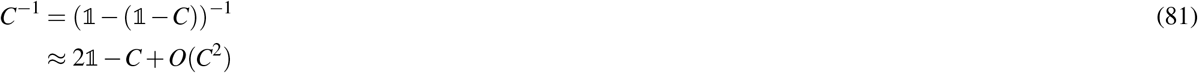

Plugging this expression back into Eqn. 71 gives us

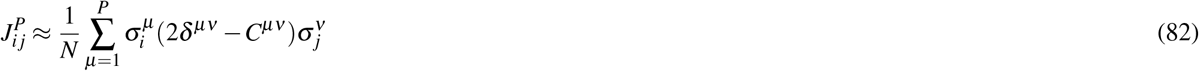

which is identical to the expression for 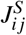 given in the last line of Eqn. 80 up to terms of 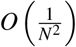.

The expansion of *C* in Eqn. 81 converges if the eigenvalues of *C* are all contained in the interval [0, 2]. For random memories with components 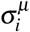 drawn iid from *𝒩* (0, 1), the distribution of the eigenvalues of *C*^*μν*^ is given by the Marchenko-Pastur distribution given by, which talks the following form for *P, N* → ∞

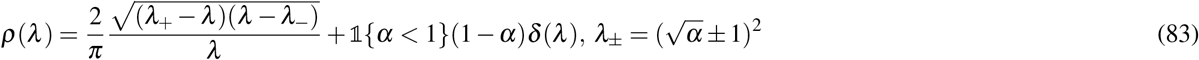

Since *λ* ∈ [*λ*_−_, *λ*_+_], the expansion of *C* is valid for *α* < 0.17 implying that 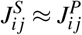 for sufficiently small *α* and large *N*.

In Fig. 6 we compare *m* as a function of *m*_0_ for two values of *α* for the three learning rules and in Fig. 7, we show *m*_*min*_ (as defined in the main text) as a function of *α*.

#### 3.5 Connection to Storage of Knowledge Structures

While the various learning rules and dynamics have quantitative effects on the values on the various network order parameters at fixed memory load and network size, their qualitative behavior as a function of the retrieval cue overlap *m*_0_ remains roughly the same for small values of *α* below capacity.

In general, we have found that outside of the basin of attraction, ⟨*m* ⟩ (and *m*\*) scales linearly with *m*_0_ for all three learning rules and takes the form in Eqn. 7 where the coefficient *f* (*α*) depends on the learning rule.

**Figure 6.**
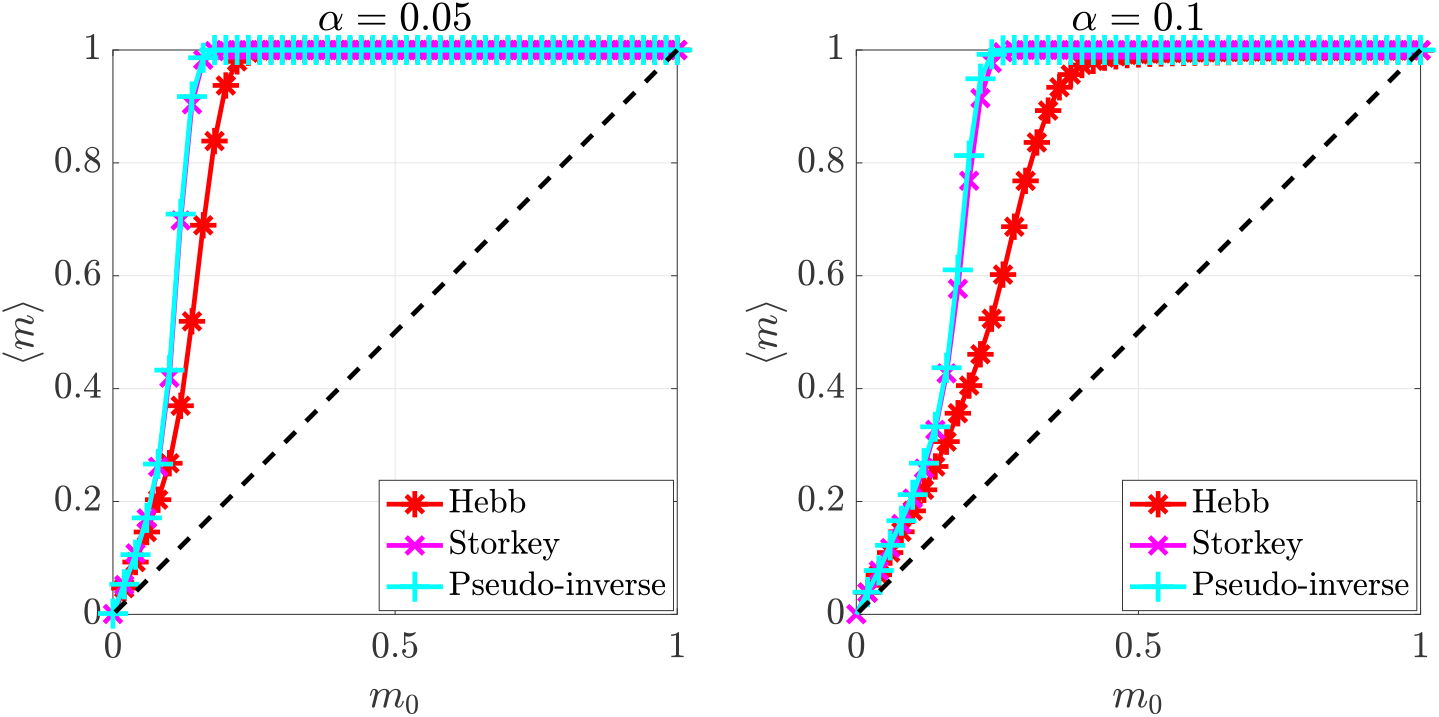
Comparison of ⟨*m* ⟩ as function of retrieval cue overlap *m*_0_ for the Hebb, Storkey, and Pseudo-inverse learning rules for two values of *α. T* = 20 parallel updates, *N* = 1000 and 50 trials are used for both figures.

**Figure 7.**
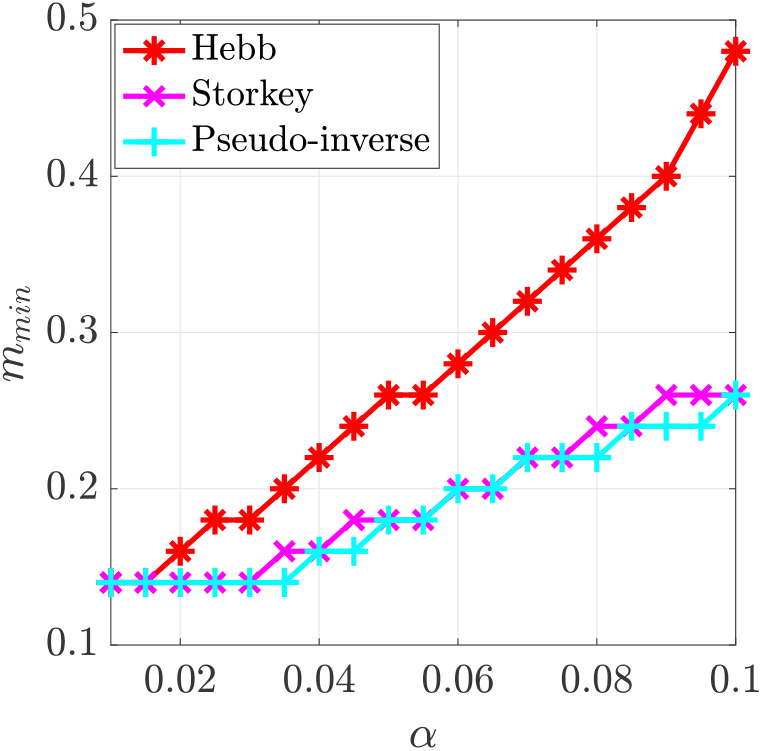
The average size of the basins of attraction (as defined in the main text) for the Hebb Storkey and Pseudo-inverse learning rules are shown together for small values of *α*.

In all cases, for large *N* where ⟨*m* ⟩ → *m*\*, the SNR behaves as

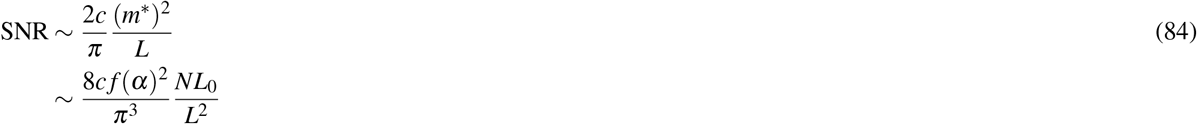

As a result, while the choice of learning rule and retrieval dynamics modify the value of *f* (*α*) defined in Eqn. 65, they do not change the scaling behavior in Eqn. 85.

Comparing *P*_*ε*_ for structures stored using the Hebb, Storkey, and Pseudo-inverse rules in Fig. 8a. In general, we find that for low values of *α* ≲ 0.3, the error of decoding from memories stored via the Storkey rule is very similar to the Pseudo-inverse rule. Both are significantly lower the error obtained when decoding from memories stored using the Hebb rule. This suggests that the Storkey learning rule sufficiently decorrelates the different structures to allow for both efficient storage and retrieval of structured knowledge.

**Figure 8.**
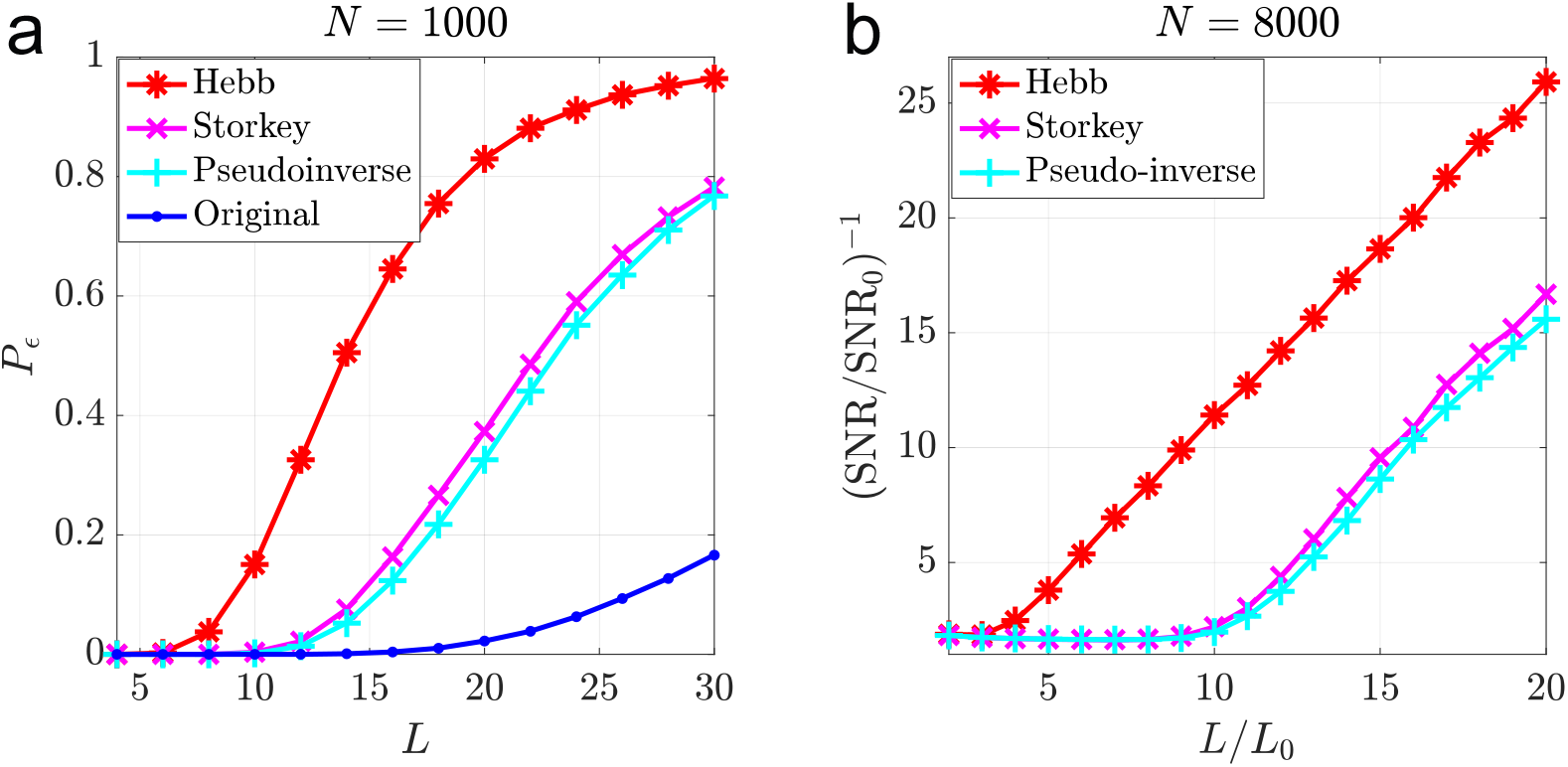
(a) Comparison of the decoding error *P*_*ε*_ for structures of size *N* = 1000 stored using the Hebb rule, Storkey rule, and Pseudo-inverse rule at memory load *α* = 0.1 and retrieved from memory with a partial cue of length *L*_0_ = 2. The *P*_*ε*_ for the original structure is also shown for comparison and the size of the dictionary is fixed to *D* = 30*N*. (b) (SNR/SNR_0_)^−1^v. *L/L*_0_ shown for *N* = 8000. *T* = 20 parallel updates are used for both figures and the is average is performed over 20, 000 memories.

### 4 Temporal Sequences as Sequences of Attractors

Previously in^7,8^, it was shown that a temporal sequence of memories could be stored in a Hopfield network by adding a second asymmetric synaptic interaction of the form

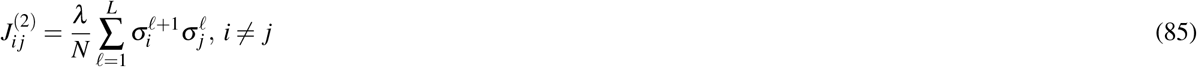

to the synaptic weight matrix, where *L* < *P* and *λ* > 1.

We can store *Q* sequences in the network by summing together a contribution 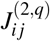 for each sequence, i.e.,

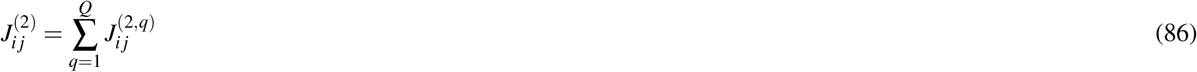

as long as each memory is only contained in one sequence.

To retrieve a sequence, one starts with an initial state *σ*_0_ = *σ* (0) within the basin of attraction of the first memory in the sequence. The evolution of the network state is then given by

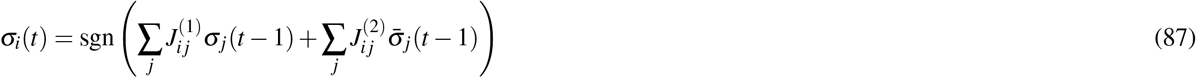

where 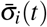 is defined as

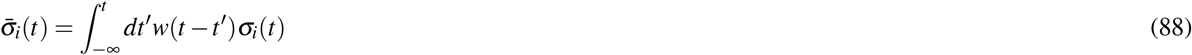

where the function *w*(*t*) represents a dynamic memory characterized by time constant *τ*. A simple choice for *w*(*t*) uses the heaviside function, i.e. 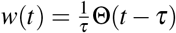, which we use for the simulation results shown in Fig. 6d of the main text. 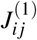 is given by a learning rule for the conventional Hopfield network such as the Hebb, Storkey, or Pseudo-inverse rules.

For long sequences, transitions between attractors will be equally spaced with period *t*_0_ after initial transients and the system will be in the pattern *μ* in the time interval ((*μ* − 1)*t*_0_, *μt*_0_). For a heaviside function *t*_0_ is given by

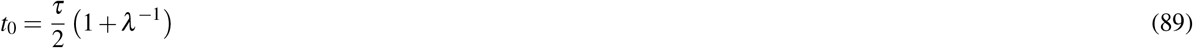

We can access the performance of this sequence memory by looking at *m*^*μ*^ (*t*) given in Eqn. 60 for each pattern in a sequence. Good performance requires *m*^*μ*^ (*t*) ≈ 1 during the interval *t* ∈ ((*μ* − 1)*t*_0_, *μt*_0_) and *m*^*μ*^ (*t*) ≪ 1 for all other *t*.

Note that if a we store two sequences sharing items in the same memory network, transitions between attractors in the dynamics given in Eqn. 87 are no longer unique.

## References

1. Tulving, E. Episodic and semantic memory. In Tulving, E. & Donaldson, W. (eds.) Organization of memory, 381–403 (Academic Press, 1972).

2. Howard, M. W. & Kahana, M. J. A distributed representation of temporal context. J. Math. Psychol. 46, 269–299 (2002).

3. Tolman, E. C. Cognitive maps in rats and men. Psychol. Rev. 55, 189–208 (1948).

4. Manns, J. R. & Eichenbaum, H. A cognitive map for object memory in the hippocampus. Learn. Mem. 16, 616–624 (2009).

5. Behrens, T. E. et al. What Is a Cognitive Map? Organizing Knowledge for Flexible Behavior. Neuron 100, 490–509 (2018).

6. Smolensky, P. Tensor product variable binding and the representation of symbolic structures in connectionist systems. Artif. Intell. 46, 159 – 216 (1990).

7. Plate, T. A. Distributed representations and nested compositional structure. (1994).

8. Eliasmith, C. How to Build a Brain: A Neural Architecture for Biological Cognition (Oxford University Press, 2013).

9. Feldman, J. & Ballard, D. Connectionist models and their properties. Cogn. Sci. 6, 205–254 (1982).

10. Holyoak, K. J. & Hummel, J. E. The proper treatment of symbols in a connectionist architecture. In Cognitive dynamics: Conceptual and representational change in humans and machines., 229–263 (Lawrence Erlbaum Associates Publishers, Mahwah, NJ, US, 2000).

11. Smolensky, P. On the Proper Treatment of Connectionism, 145–206 (Springer Netherlands, Dordrecht, 1990).

12. Feldman, J. The neural binding problem(s). Cogn. Neurodynamics 7, 1–11 (2013).

13. Jackendorff, R. Foundations of language: Brain, meaning, grammar, evolution (Oxford University Press, 2002).

14. Greff, K., van Steenkiste, S. & Schmidhuber, J. On the Binding Problem in Artificial Neural Networks. arXiv 2012.05208 (2020).

15. Barak, O., Rigotti, M. & Fusi, S. The sparseness of mixed selectivity neurons controls the generalization-discrimination trade-off. J. Neurosci. 33, 3844–3856 (2013).

16. Podlaski, W. F., Agnes, E. J. & Vogels, T. P. Context-modular memory networks support high-capacity, flexible, and robust associative memories. bioRxiv 2020.01.08.898528 (2020).

17. Kanerva, P. Binary spatter-coding of ordered k-tuples. In Artificial Neural Networks - ICANN 96, 1996 International Conference, Bochum, Germany, July 16-19, 1996, Proceedings, 869–873 (1996).

18. Levy, S. D. & Gayler, R. Vector symbolic architectures: A new building material for artificial general intelligence. In Proceedings of the 2008 Conference on Artificial General Intelligence 2008: Proceedings of the First AGI Conference, 414–418 (IOS Press, NLD, 2008).

19. Rachkovskij, D. A. & Kussul, E. M. Binding and normalization of binary sparse distributed representations by context-dependent thinning. Neural Comput. 13, 411–452 (2001).

20. Kleyko, D., Rachkovskij, D. A., Osipov, E. & Rahimi, A. A survey on hyperdimensional computing aka vector symbolic architectures, part ii: Applications, cognitive models, and challenges. ACM Comput. Surv. (2022).

21. Stewart, T. C., Bekolay, T. & Eliasmith, C. Neural representations of compositional structures: representing and manipulating vector spaces with spiking neurons. Connect. Sci. 23, 145–153 (2011).

22. Schlegel, K., Neubert, P. & Protzel, P. A comparison of vector symbolic architectures. 0123456789 (Springer Netherlands, 2021).

23. Battaglia, P. W. et al. Relational inductive biases, deep learning, and graph networks. arXiv 1806.01261 [cs.LG] (2018).

24. Santoro, A. et al. A simple neural network module for relational reasoning. In Guyon, I.et al. (eds.) Advances in Neural Information Processing Systems, vol. 30 (Curran Associates, Inc., 2017).

25. Zaheer, M. et al. Deep sets. In Guyon, I.et al. (eds.) Advances in Neural Information Processing Systems 30, 3391–3401 (Curran Associates, Inc., 2017).

26. Devlin, J., Chang, M. W., Lee, K. & Toutanova, K. BERT: Pre-training of deep bidirectional transformers for language understanding. NAACL HLT 2019 - 2019 Conf. North Am. Chapter Assoc. for Comput. Linguist. Hum. Lang. Technol. - Proc. Conf. 1, 4171–4186 (2019).

27. Frady, E. P., Kleyko, D. & Sommer, F. T. A Theory of Sequence Indexing and Working Memory in Recurrent Neural Networks. Neural Comput. 30, 1449–1513 (2018).

28. Whittington, J. C. et al. The Tolman-Eichenbaum Machine: Unifying Space and Relational Memory through Generalization in the Hippocampal Formation. Cell 183, 1249–1263.e23 (2020).

29. Whittington, J. C. R., Warren, J. & Behrens, T. E. J. Relating transformers to models and neural representations of the hippocampal formation. In ICLR (2022).

30. Gemici, M. et al. Generative temporal models with memory. arXiv 1702.04649 [cs.LG] (2017).

31. Cox, G. E. & Criss, A. H. Parallel interactive retrieval of item and associative information from event memory. Cogn. Psychol. 97, 31–61 (2017).

32. Amit, D. J., Gutfreund, H. & Sompolinsky, H. Storing infinite numbers of patterns in a spin-glass model of neural networks. Phys. Rev. Lett. 55, 1530–1533 (1985).

33. Hopfield, J. J. Neural networks and physical systems with emergent collective computational abilities. Proc. Natl. Acad. Sci. 79, 2554–2558 (1982).

34. O’Reilly, R. C. & McClelland, J. L. Hippocampal conjunctive encoding, storage, and recall: Avoiding a trade-off. Hippocampus 4, 661–682 (1994).

35. Sompolinsky, H. & Kanter, I. Temporal association in asymmetric neural networks. Phys. Rev. Lett. 57, 2861–2864 (1986).

36. Xie, X., Hahnloser, R. & Seung, H. S. Groups of Neurons in Lateral Inhibitory Networks..

37. Tsodyks, M. V. & Feigel’man, M. V. The enhanced storage capacity in neural networks with low activity level. Europhys. Lett. (EPL) 6, 101–105 (1988).

38. Amit, D. J., Gutfreund, H. & Sompolinsky, H. Spin-glass models of neural networks. Phys. Rev. A 32, 1007–1018 (1985).

39. Kanter, I. & Sompolinsky, H. Associative recall of memory without errors. Phys. Rev. A 35, 380–392 (1987).

40. Amit, D. J., Gutfreund, H. & Sompolinsky, H. Statistical mechanics of neural networks near saturation. Annals Phys. 173, 30 – 67 (1987).

41. Logan, G. D. Automatic control: How experts act without thinking. Psychol. Rev. 125, 453–485 (2018).

42. Logan, G. D. & Cox, G. E. Serial memory: Putting chains and position codes in context. Psychol. review 128, 1197–1205 (2021).

43. Nickel, M., Rosasco, L. & Poggio, T. Holographic embeddings of knowledge graphs. In Proceedings of the Thirtieth AAAI Conference on Artificial Intelligence, AAAI’16, 1955–1961 (AAAI Press, 2016).

44. Silver, R. A. Neuronal arithmetic. Nat. Rev. Neurosci. 11, 474–489 (2010).

45. Mehaffey, W. H., Doiron, B., Maler, L. & Turner, R. W. Deterministic multiplicative gain control with active dendrites. J. Neurosci. 25, 9968–9977 (2005).

46. Ahmadian, Y., Rubin, D. B. & Miller, K. D. Analysis of the Stabilized Supralinear Network. Neural Comput. 25, 1994–2037 (2013).

47. Deng, P. Y. & Klyachko, V. A. The diverse functions of short-term plasticity components in synaptic computations. Commun. Integr. Biol. 4, 543–548 (2011).

48. Zucker, R. S. & Regehr, W. G. Short-term synaptic plasticity. Annu. Rev. Physiol. 64, 355–405 (2002).

49. Ba, J. et al. [2016-NIPS] Using Fast Weights to Attend to the Recent Past. 1–9 (2016).

50. Schlag, I., Irie, K. & Schmidhuber, J. Linear transformers are secretly fast weight programmers. In ICML (2021).

51. Frady, E. P., Kleyko, D. & Sommer, F. T. Variable Binding for Sparse Distributed Representations: Theory and Applications. IEEE Transactions on Neural Networks Learn. Syst. 1–14 (2021).

52. Rachkovskij, D. A. Representation and processing of structures with binary sparse distributed codes. IEEE Transactions on Knowl. Data Eng. 13, 261–276 (2001).

53. Rachkovskij, D. A., Kussul, E. M. & Baidyk, T. N. Building a world model with structure-sensitive sparse binary distributed representations. Biol. Inspired Cogn. Archit. 3, 64–86 (2013).

54. Hiratani, N. & Sompolinsky, H. Optimal quadratic binding for relational reasoning in vector symbolic neural architectures. arXiv 2204.07186 [q–bio.NC] (2022).

55. Franklin, N. T., Norman, K. A., Ranganath, C., Zacks, J. M. & Gershman, S. J. Structured Event Memory: A neuro-symbolic model of event cognition. Psychol. Rev. 127, 327–361 (2020).

56. Cox, G. E. & Criss, A. H. Similarity leads to correlated processing: A dynamic model of encoding and recognition of episodic associations. Psychol. Rev. 102, 792–828 (2020).

57. Zeng, T., Tompary, A., Schapiro, A. C. & Thompson-Schill, S. L. Tracking the relation between gist and item memory over the course of long-term memory consolidation. eLife 10, e65588 (2021).

58. Cox, G. E. & Shifrin, R. M. A dynamic approach to recognition memory. Psychol. Rev. 124, 795–860 (2017).

59. Kumar, A. A. Semantic memory: A review of methods, models, and current challenges. Psychon. Bull. Rev. 28, 40–80 (2021).

60. McClelland, J. L., McNaughton, B. L. & O’Reilly, R. C. Why there are complementary learning systems in the hippocampus and neocortex: Insights from the successes and failures of connectionist models of learning and memory. Psychol. Rev. 102, 419–457 (1995).

61. Sun, W., Advani, M., Spruston, N., Saxe, A. & Fitzgerald, J. E. Organizing memories for generalization in complementary learning systems. bioRxiv 2021.10.13.463791 (2021).

62. Plate, T. A. Holographic reduced representations. IEEE Transactions on Neural Networks 6, 623–641 (1995).

63. Storkey, A. Increasing the capacity of a hopfield network without sacrificing functionality. In Proceedings of the 7th International Conference on Artificial Neural Networks, ICANN ‘97, 451–456 (Springer-Verlag, Berlin, Heidelberg, 1997).

64. Storkey, A. & Valabregue, R. A hopfield learning rule with high capacity storage of time-correlated patterns (1997).

## References

1. Plate, T. A. Distributed representations and nested compositional structure. (1994).

2. Kanter, I. & Sompolinsky, H. Associative recall of memory without errors. Phys. Rev. A 35, 380–392 (1987).

3. Storkey, A. Increasing the capacity of a hopfield network without sacrificing functionality. In Proceedings of the 7th International Conference on Artificial Neural Networks, ICANN ‘97, 451–456 (Springer-Verlag, Berlin, Heidelberg, 1997).

4. Storkey, A. & Valabregue, R. A hopfield learning rule with high capacity storage of time-correlated patterns (1997).

5. Amit, D. J., Gutfreund, H. & Sompolinsky, H. Storing infinite numbers of patterns in a spin-glass model of neural networks. Phys. Rev. Lett. 55, 1530–1533 (1985).

6. Storkey, A. J. Efficient Covariance Matrix Methods for Bayesian Gaussian Processes and Hopfield Neural Networks. 138 (1999).

7. Sompolinsky, H. & Kanter, I. Temporal association in asymmetric neural networks. Phys. Rev. Lett. 57, 2861–2864 (1986).

8. Kleinfeld, D. & Sompolinsky, H. Associative neural network model for the generation of temporal patterns. Theory and application to central pattern generators. Biophys. J. 54, 1039–1051 (1988).

